# Suboptimal human inference inverts the bias-variance trade-off for decisions with asymmetric evidence

**DOI:** 10.1101/2020.12.06.413591

**Authors:** Tahra L Eissa, Joshua I Gold, Krešimir Josić, Zachary P Kilpatrick

**Author notes:** equal contributions (Joshua I Gold, Krešimir Josić, Zachary P Kilpatrick).

## Abstract

Solutions to challenging inference problems are often subject to a fundamental trade-off between bias (being systematically wrong) that is minimized with complex inference strategies and variance (being oversensitive to uncertain observations) that is minimized with simple inference strategies. However, this trade-off is based on the assumption that the strategies being considered are optimal for their given complexity and thus has unclear relevance to the frequently suboptimal inference strategies used by humans. We examined inference problems involving rare, asymmetrically available evidence, which a large population of human subjects solved using a diverse set of strategies that were suboptimal relative to the Bayesian ideal observer. These suboptimal strategies reflected an inversion of the classic bias-variance trade-off: subjects who used more complex, but imperfect, Bayesian-like strategies tended to have lower variance but high bias because of incorrect tuning to latent task features, whereas subjects who used simpler heuristic strategies tended to have higher variance because they operated more directly on the observed samples but displayed weaker, near-normative bias. Our results yield new insights into the principles that govern individual differences in behavior that depends on rare-event inference, and, more generally, about the information-processing trade-offs that are sensitive to not just the complexity, but also the optimality of the inference process.

## Introduction

Understanding how the brain makes inferences about the world requires first understanding the diversity of strategies individuals use to solve inference problems. One useful approach for understanding this diversity is to assess patterns of errors, which can reflect particular inference strategies. In general, errors can result from either: 1) bias, which can arise due to an incorrect model of the world that gives rise to inferences that are systematically offset from the ground truth; or 2) variability, which can reflect either intrinsic noise or oversensitivity to changes in observations (which we refer to as ”noise” and ”variance”, respectively), either of which can lead to inferences that are variable over multiple instances of the same problem. Some forms of human inference reflect an inherent trade-off between bias and variance [1] that depends on the complexity of the inference process [2, 3]: higher complexity provides more flexibility that tends to decrease bias but at the expense of oversensitivity to noise, whereas lower complexity tends to increase bias but decrease variance. However, this trade-off has typically been considered in the context of inference processes (or “models” in machine learning) that vary in complexity but, for a given complexity, are optimized for the given problem. Much less understood is whether and how similar trade-offs can be used to understand the diverse forms of suboptimal strategies that people often use to solve inference problems [4–6].

To better understand sources of errors in suboptimal inference, and how they might relate to the bias-variance trade-off, we examined the choice behavior of human subjects performing a two-alternative forced-choice task for which evidence in favor of one choice was sparse across both conditions [7]. These inference problems are interesting because, as we detail below, they give rise to choice asymmetries, i.e., a tendency to chose one alternative more frequently than the other, even when the alternatives are *a priori* equally likely. We exploited this tendency to identify how the subjects’ choice strategies differed in terms of their: 1) bias, defined as the choice asymmetry relative to that of an ideal observer; and 2) variance, defined as choice variability relative to that of an ideal observer. We were particularly interested in how these behavioral patterns differed across individual subjects and task conditions, and how they related to different models that we fit to behavior.

We focused on two classes of models whose differences were central to the interpretation of our primary findings. The first was based on Bayesian principles. Models in this class differed in their exact implementation but depended on one or more latent variables that captured key task features that drove choice asymmetries. The second class contained models based on simpler, heuristic principles which more directly mapped patterns of observations to choices. Subjects using Bayesian-like strategies ranged from using nearly optimal versions, with low bias and variance, to suboptimal versions that tended to have higher bias but low variance. In contrast, subjects using heuristic strategies tended to have higher variance but low bias. This inverted form of bias-variance trade-off followed from the differences in the two strategies: suboptimalites in Bayesian-like strategies involved mistuned estimates of task-relevant latent variables (i.e., being wrong about their true values), which led to excess choice biases without adding much choice variability, whereas suboptimalities in heuristic strategies involved operating more directly on specific sequences of observations, which led to increased variance without causing systematic biases. These results highlight how breaking the symmetry of standard decision tasks and considering the diversity of strategies used by individuals can reveal systematic relationships between strategy complexity, bias-variance balance, and the sources of human errors.

## Results

We used a task based on a classic inference problem that required each subject to infer which of two *a priori* equally likely jars filled with red and blue balls was the source of a sample of balls drawn with replacement (Fig. 1**a**). On each trial, the sample of 2, 5, or 10 balls was shown all at once, with the contents of each jar visible at all times. Subjects were told that each jar was equally likely to be the source (Fig. S1). Across different blocks, the proportions of red and blue balls in each jar were varied, thereby altering the ideal evidence weight of each observation. Under “symmetric” conditions, the ratios of the two ball colors in the two jars were reciprocal. In contrast, under “asymmetric” conditions, the ratios were non-reciprocal. Specifically, one ball color (either blue or red) accounted for fewer than half the balls in both jars and thus was “rare”. This rare color was selected randomly for each subject but was then fixed throughout the experiment. Thus, the jar with more rare balls was termed the “high” jar, and the jar with fewer rare balls was termed the “low” jar. We asked how ideal Bayesian observers and human observers compare in their use of symmetric and asymmetric information to infer which jar is the source of an observed sample.

**Figure 1:**
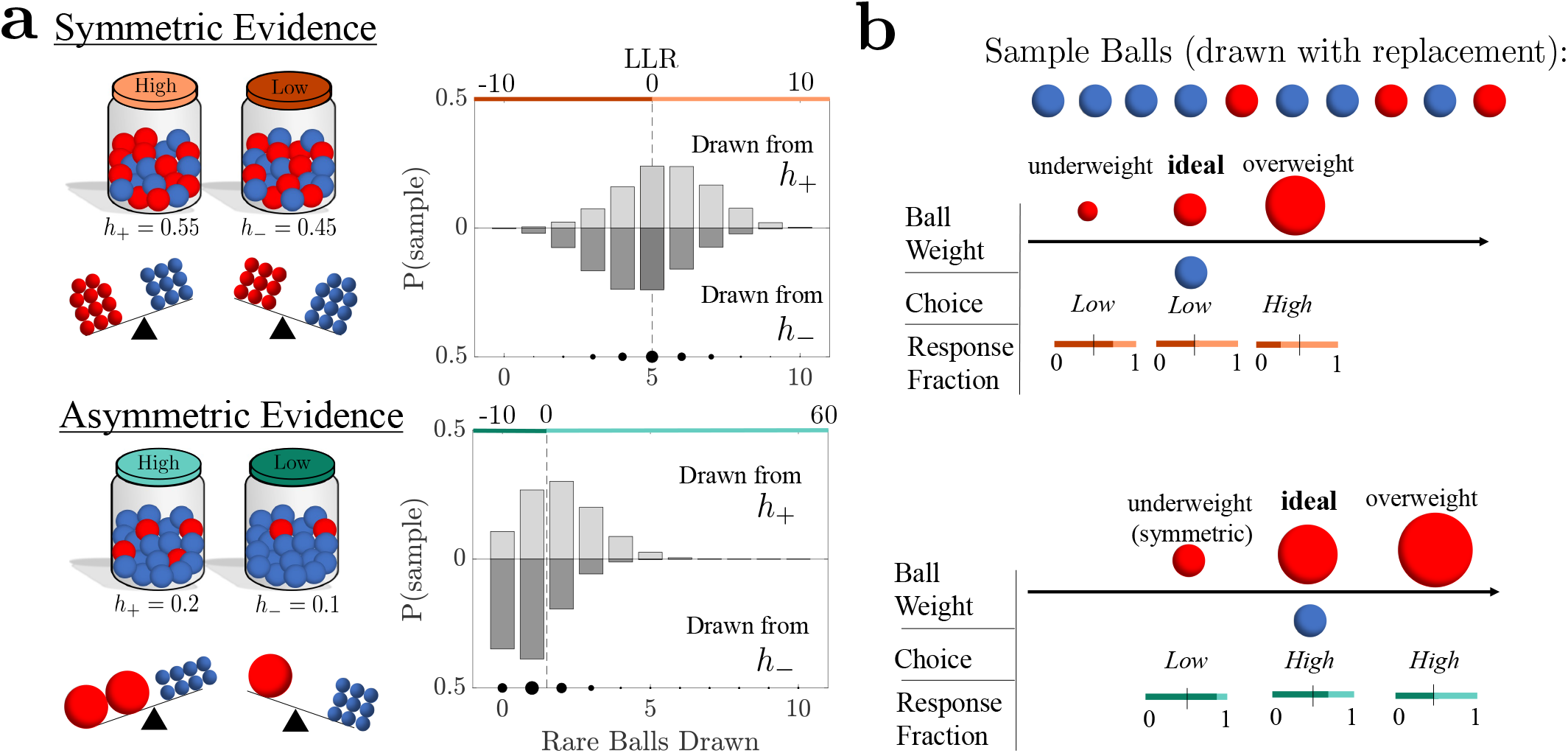
Different environmental evidence weights cause decision biases. **a**. Schematic of the Jar-Discrimination Task. Balls were drawn with replacement from one of two equally probable jars with different ratios of red to blue balls. Here *h*_*±*_ denotes the probability that a red ball is drawn from the high (*h*_+_) and low (*h*_*−*_) jar. We considered conditions with symmetric priors and symmetric evidence (*h*_*−*_ = 1 *− h*_+_), in which the weight of red/blue balls had equal amplitude but opposite sign, or asymmetric evidence (*h*_*−*_ ≠ 1*− h*_+_), in which rare (in this example red) balls were weighted more heavily in a decision. Right panels depict the corresponding probability distribution of a 10-ball sample for a given number of rare balls for the high jar (*h*_+_, top) and low jar (*h*_*−*_, bottom). Colored bars denote an ideal Bayesian observer’s jar choice resulting from log likelihood ratio (LLR) presented on the top axis (an LLR of zero results in a random response). **b**. Example of a 10-ball sample and corresponding choices of a Bayesian observer with varying relative ball weights. Top: Ideal ball weights for the symmetric environment produce even response fractions. Bottom: Ideal asymmetric weights produce a choice asymmetry in favor of the low jar. Deviations from the ideal weights in either environment produce decision biases.

### Ideal Bayesian Observer

We first derived the strategy of an ideal Bayesian observer that knows the task structure. The parameter *h*_*±*_ refers to the proportion of rare-colored balls in the given jar: The *h*_+_ (high) jar included more balls of the rare color, whereas the *h*_*−*_ (low) jar included fewer balls of the rare color, so that 0 *< h*_*−*_ *< h*_+_. When the proportions were symmetric, *h*_+_ = 1 *− h*_*−*_. When the proportions were asymmetric, 0 *< h*_*−*_ *< h*_+_ *<* 0.5 (Fig. 1**a**). Because the two jars were always visible, we assumed the fractions of rare balls, *h*_+_ and *h*_*−*_, in the low and high jars are known to the ideal observer.

The ideal observer sees a sample of draws, *ξ*_1:*n*_, where *ξ*_*i*_ = 1(*ξ*_*i*_ = *−*1) if a rare (common) ball is drawn, and computes as evidence the log-odds ratio (*belief*), 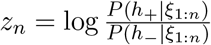, between the probabilities that the sample of draws came from either jar. Balls were seen all at once, but the recursive belief-update equation shows the effect of each observation,

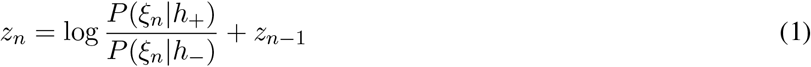

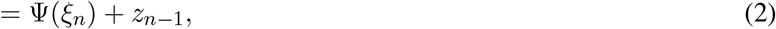

where the belief increment after the *n*^th^ observation is

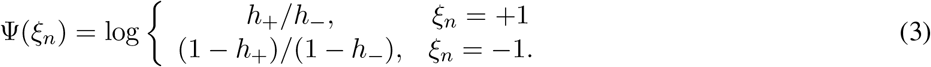

The most likely alternative given *n* observations is determined by the sign of *z*_*n*_: *z*_*n*_ *>* 0 → choose the high jar; *z*_*n*_ *<* 0 → choose the low jar. On symmetric trials, the ratio of ball colors is reciprocal, and thus the magnitude ofthe belief increments is the same for either observation, 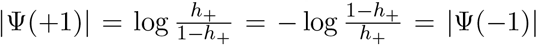. In contrast, on asymmetric trials (*h*_*−*_ ≠ 1 *− h*_+_), the magnitudes of the belief updates are not equal. Thus, for rare asymmetric evidence, 0 *< h*_*−*_ *< h*_+_ *<* 0.5, observing a rare ball leads to a larger belief update than observing a common ball (| Ψ(+1) | *>* |Ψ(−1) |) (Fig. 1**a**, bottom schematics).

The impact of evidence asymmetry on ideal-observer choices can be illustrated by comparing the probability distributions of rare balls in a 10-ball sample (Fig. 1**a**, right plots). For symmetric jars, the distributions of rare-ball counts for the low and high jars, and the evidence (LLR) they provide, are symmetric about the midline at 5 observed rare balls. Thus, the ideal observer’s beliefs and choices are also symmetric in this environment, and in line with the prior (Fig. 1**b**, top). In contrast, for asymmetric jars the distributions of rare-ball counts are not reflections of one another for the two jars and are skewed based on the *h* values. For the asymmetric jars shown, counts of zero or one rare ball(s), corresponding to evidence favoring the low jar, occur more often than counts of two or more rare balls, corresponding to evidence favoring the high jar. This is true across both jars (chosen at random). Thus, in the asymmetric case, even the use of the correct, symmetric prior and the appropriate weighting of evidence leads to a choice asymmetry with a larger expected fraction of low-jar choices (Fig. 1**b**, bottom).

In principle, suboptimal inference with asymmetric evidence could take several forms (Figs. 1, 2**a**). For example, there could be systematic choice biases, which might result from: 1) mistuning the evidence weight (LLR), for instance by underweighting or overweighting rare balls, which would, respectively, increase or decrease the choice asymmetries exhibited by the ideal observer; and/or 2) mistuning the prior, which occurs when the observer’s prior does not match the true prior. In addition to these effects, internal noise, heuristic strategy components not based on the LLR, or other factors could lead to choice variability.

**Figure 2:**
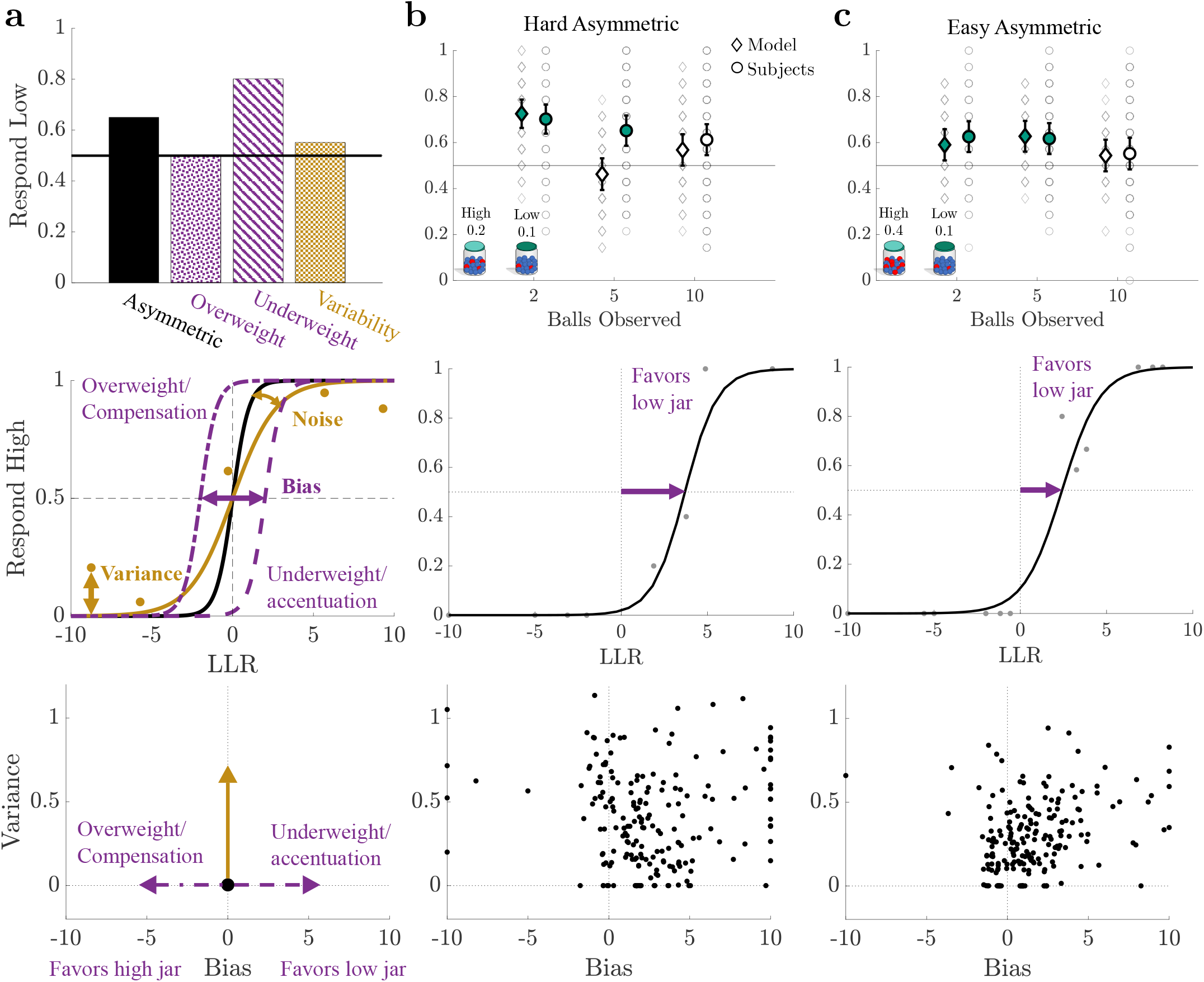
Ideal observers and human participants show similar behaviors, on average. **a**. Top: A mistuning of ball weights decreases (overweighting) or increases (underweighting) choice asymmetry in environments with asymmetric evidence, whereas increases in variability (inclusion of noise and/or variance) cause minimal impact on choice asymmetry. Center: An increase in noise decreases the slope, and a bias results in a horizontal shift of the psychometric function. We define variance as the mean absolute error between the fit psychometric function and the data. Bottom: Schematic of bias and variance effects as shown in the data figures to the right. Bias was bounded between [***−*** 10, 10] to mitigate overfitting due to outliers. Positive (negative) biases corresponded to more (fewer) low-jar selections. **b**. Individual and population results for the Hard Asymmetric block. Top: Low jar-response fractions for each subject (*N* = 201, grey circles, overlap due to discretization) and sample-matched ideal observer responses (grey diamonds) for all sample lengths (number of balls observed). Population bootstrapped means (1000 iterations) and 95% confidence intervals shown. Filled markers denote a significant population shift away from 0.5 (*p <* 0.05). Center: Summary psychometric function (line) based on the median subject responses (dots) for each block across all sample lengths. Bottom: All individual subject bias and variance parameter fits computed using logistic regression. **c**. Metrics as in **b** for the Easy Asymmetric block.

### Human behavior

We used the crowdsourcing platform Amazon Mechanical Turk (MTurk) to recruit 201 subjects to perform the Jar-Discrimination task (Fig. 1**a**). Each subject performed 60 trials in a control block (CT; *h*_+_ = 0.9*/h*_*−*_ = 0.1) to acclimate to the task and then 42 trials under each of four conditions that varied in difficulty and evidence asymmetry: hard asymmetric (HA, *h*_+_ = 0.2*/h*_*−*_ = 0.1), hard symmetric (HS, *h*_+_ = 0.55*/h*_*−*_ = 0.45), easy asymmetric (EA, *h*_+_ = 0.4*/h*_*−*_ = 0.1), and easy symmetric (ES, *h*_+_ = 0.7*/h*_*−*_ = 0.3). The task structure and parameters were informed by simulations and pilot studies. Some of the analyses we describe below were pre-registered (see Methods for a detailed breakdown). Below we focus on results from the asymmetric conditions and provide results from the symmetric conditions in the Supplemental Materials for comparison purposes.

On average, the subjects tended to make choices that were roughly consistent with predictions of the ideal observer (Fig. S2). In particular, for asymmetric conditions both the ideal observer and the subjects tended to have choice asymmetries that were strongest on shorter (two- or five-ball) trials (Fig. 2**b,c**, top). These choice asymmetries were reduced when trials that included one or more rare balls (rare-ball trials) were more common (Fig. S3), and they did not depend on the specific sequence of the sampled balls (Fig. S4). For symmetric conditions, both the ideal observer and the subjects tended to make similar fractions of low/high choices (Fig. S5).

To identify potential suboptimalities in the behavior of individual subjects, particularly in the asymmetric conditions, we fit each subject’s choice data in each block to a logistic psychometric function describing the fraction of high-jar choices as a function of the LLR in favor of the high jar (Fig. 2**a**, middle). For an ideal observer, the psychometric function is a step function (with no choice variability except random guessing on LLR=0 trials) with no bias (i.e., the step is at LLR=0; note that, as shown in Fig. 1, this unbiased psychometric function can nonetheless corresponds to asymmetric choice fractions when the evidence more often favors the low jar). For subject data, we defined bias as the horizontal shift of the psychometric function. We decomposed choice variability into two components defined in terms of: 1) the inverse slope of the best-fitting function, which we interpret primarily as noise because it represents choice variability (uncertainty) that is associated with LLR; and 2) the mean absolute error between the subject data and the predicted psychometric function, which likely includes noise but also variance because it represents choice patterns that are disassociated from the LLR. That is, this form of choice variability represents choice patterns that deviate from the smooth, LLR-based psychometric function because they occur in response to particular, observed ball samples, even if samples share the same LLR (see Methods for more details). In the main text, below, we focus on variance but include comparable analyses of noise in the supplementary figures to show that our conclusions are consistent with both measures of choice variability. These metrics allowed us to distinguish the impact of bias and choice variability individually on each subject’s performance (Fig. 2**a**, bottom).

Individual subjects exhibited a range of behavioral suboptimalities, including errors attributable to bias, variance, and/or noise that varied in magnitude across subjects (Figs. 2**b,c** middle and bottom plots and S6). Variance ranged from values of zero, corresponding to choices that exactly matched the best-fitting psychometric function, to near one, corresponding to choice patterns that deviated substantially from the smooth psychometric function (Fig. S6). The slopes of the psychometric functions, corresponding to the inverse of noise, ranged from infinitely steep to zero across subjects and task conditions (Fig. S7). Bias also varied in magnitude but tended in the same direction across subjects, in most cases corresponding to an increase in the choice asymmetries favoring the low jar. These biases occurred throughout each block and thus did not appear to involve a learning mechanism that might have compensated for a mismatch between the known, symmetric prior and a growing understanding through each block that the choice fractions did not respect that symmetry (Fig. S8). Instead, these biases were consistent with a process of evidence underweighting or choice accentuation, as described above. Below we analyze these different kinds of errors in more detail by considering the specific strategies that might have produced them.

### Formal comparison of Bayesian and heuristic models

To relate these human behavioral patterns to particular inference strategies, we fit choice data from each subject and each block to different parameterized Bayesian-based and heuristic models (Fig. 3). We then determined the bias-variance trends for each subject’s best-fitting model.

**Figure 3:**
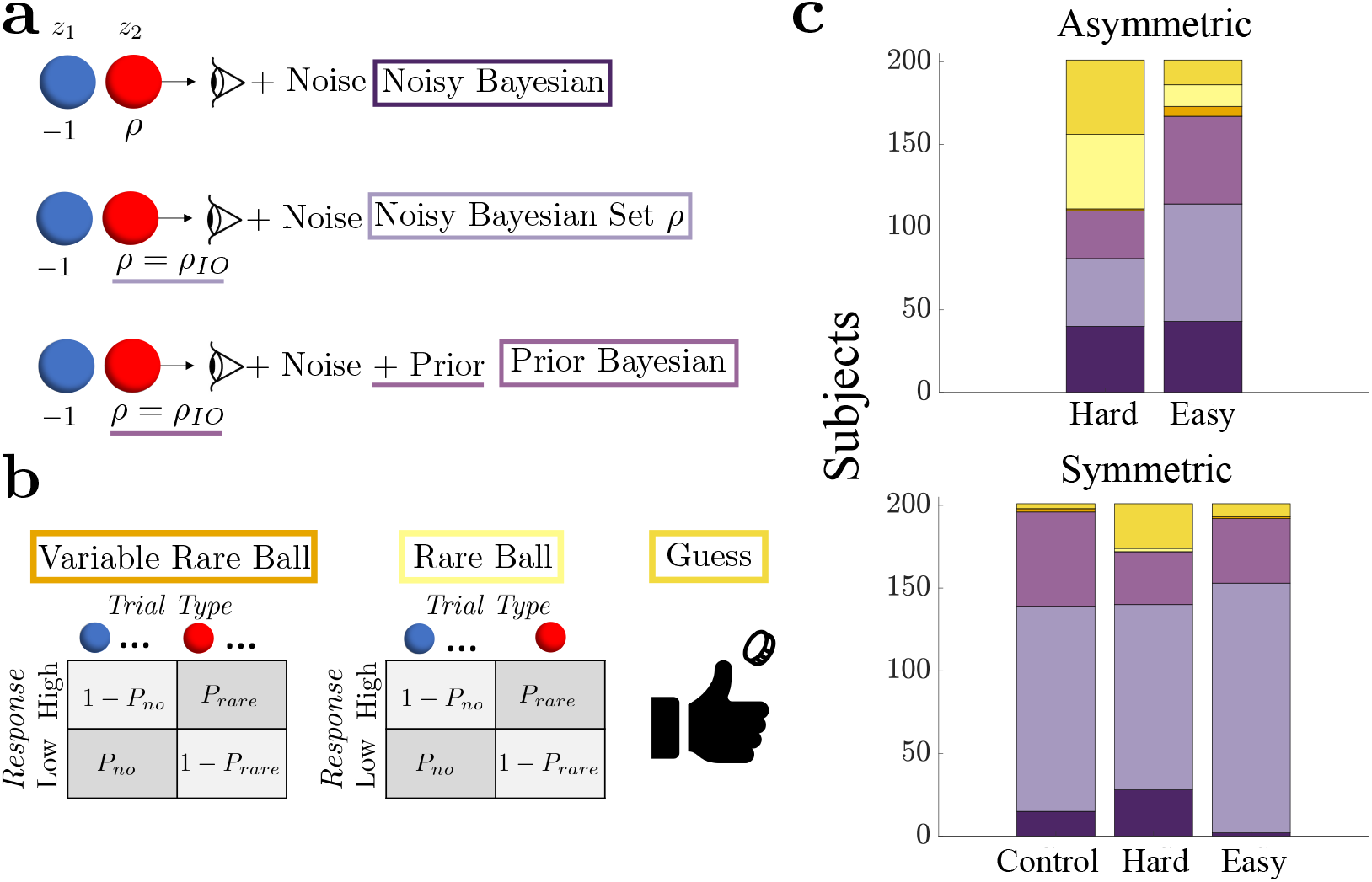
Decision-making model types and fits. **a**. Bayesian models. The Noisy Bayesian model includes noise and a rare ball weight, *ρ*, that varies across subjects. The Noisy Bayesian Set *ρ* model assumes that *ρ* equals the ideal observer’s rare-ball weight (*ρ*_*IO*_). The Prior Bayesian model includes a jar bias (prior), and assumes *ρ* = *ρ*_*IO*_. Differences from the Noisy Bayesian model are underlined. **b**. Heuristic models. The probability of a response in the Variable Rare Ball model depends on whether or not the number of observed rare balls exceeds a threshold. This threshold is inferred. The Rare Ball model is a reduction of the Variable Rare Ball model with the response probability determined by the presence or absence of one or more rare balls (threshold is set to 1). Under the Guess model, the high jar is chosen with with some probability that is set as a free parameter. **c**. Subjects categorized by the model that best matched their responses. The responses of a majority of subjects in all blocks were best described by Bayesian models (98% in CT, 55% in HA, 86% in HS, 86% in EA, and 95% in ES), but with a relatively high percentage of heuristic strategies under the HA condition.

We used three Bayesian-based models (Fig. 3**a**). The first included a noisy version of the log-likelihood,

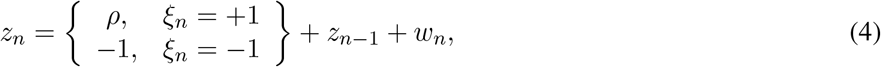

where *w*_*n*_ is a normally distributed random variable with zero mean and variance *a*^2^, and included *ρ* as a free parameter representing the belief update in response to observing a rare ball (“Noisy Bayesian”). When *ρ >* 1, the model weights a rare-ball observation more strongly than an observation of a common ball. In the second model, we set

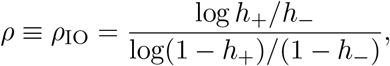

When noise is absent, this version is equivalent to the ideal observer model (“Noisy Bayesian Set *ρ*”). In the third model, we added a parameterized prior to the “Noisy Bayesian set *ρ*” model (“Prior Bayesian”).

We also modeled several heuristic strategies that were not based on likelihoods computed from samples (Fig. 3**b**). The first assumed that the probability of choosing the high jar is determined by whether the number of observed rare balls exceeded a threshold. This threshold was a model parameter whose value we inferred from subject responses (“Variable Rare Ball”). Since the threshold was fixed regardless of the total ball count (2, 5, or 10), the model could produce different response probabilities for different ball patterns with the same LLR. The second model was based on the assumption that the observer chooses the high jar with probability *P*_rare_ whenever observing one or more rare balls in a sample, and with probability 1 *− P*_no_ otherwise (“Rare Ball”). This is equivalent to fixing the threshold parameter in the first, Variable Ball model to 1. The third model described a simple guessing strategy, in which the observer selects the high jar with some probability, regardless of the observations (“Guess”).

We used Bayes factors to select the model that best matched each subject’s responses on each block separately (after confirming model identifiability; see Figs. S9, S10, and Methods). Across all blocks, most subjects exhibited choice behaviors that were most consistent with one of the Bayesian models (Figs. 3**c** and S11). For symmetric blocks, this tendency was strongest. For asymmetric blocks, particularly under the Hard Asymmetric condition, strategies were more diverse, with many subjects identified as using heuristic strategies. Individual subject’s responses did not typically match the same model across blocks, but approximately half (49%) of subjects were consistently well matched to models in the Bayesian class across blocks (Fig. S12).

### Model-dependent bias-variance trends

For the asymmetric conditions, we found a systematic relationships between an individual subject’s best-fitting strategy type and the magnitude of their bias and variance as determined by the fitted psychometric functions (Fig. 4**a**; we observed similar relationships when considering noise instead of variance, Fig. S13 bottom). Specifically, responses of subjects best fit by the nearly optimal Bayesian model (i.e., the Noisy Bayesian Set *ρ* model) were characterized by almost no bias and variance and thus had choice asymmetries that were similar to the ideal observer. However, the remaining subjects exhibited suboptimalities that differed depending on whether the responses of the subject were best fit by a heuristic or a Bayesian-like model. Subjects best fit by heuristic models had biases that exhibited substantial individual variability but were, on average, close to zero under both of the asymmetric conditions, and they tended to have relatively high variance. In contrast, subjects best fit by suboptimal Bayesian-like models (the Noisy Bayesian or Prior Bayesian model) tended to have relatively low variance but responses strongly biased towards the low jar.

**Figure 4:**
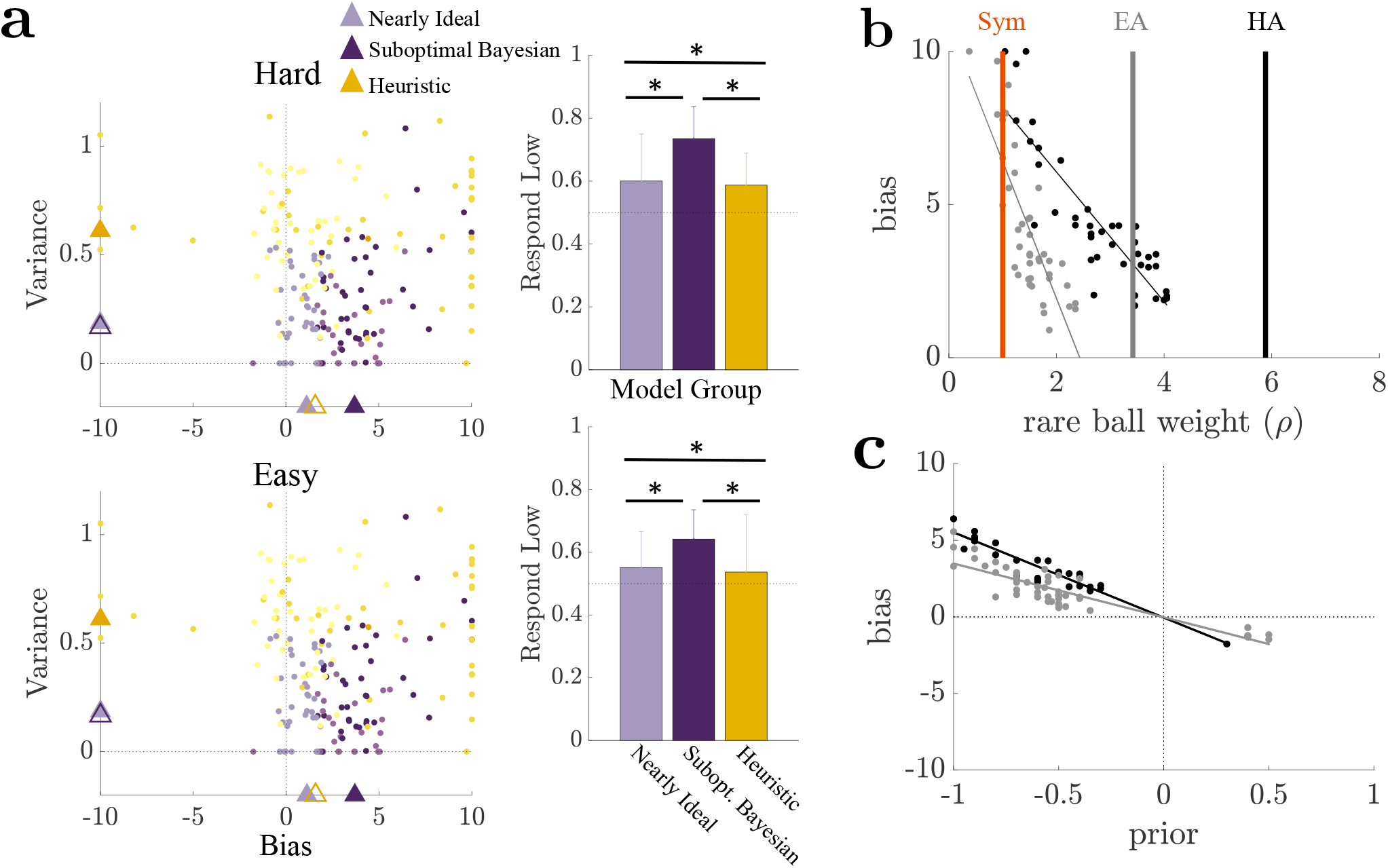
Increased bias and variance in asymmetric blocks correspond to suboptimal Bayesian and heuristic model fits, respectively. **a**. Left: Asymmetric block bias-variance plots from Fig. 2, color-coded according to each subject’s best-fitting model as in Fig. 3, and the model best describing the responses of each subject. Triangles denote subject medians for: 1) nearly ideal subjects (best fit to “Noisy Bayesian Set *ρ*” model), 2) suboptimal Bayesian subjects (best fit to “Noisy Bayesian” or “Prior Bayesian” model), 3) heuristic subjects. Statistics of suboptimal Bayesian and heuristic subjects denoted by filled (open) triangles were significantly (not significantly) different from nearly ideal subjects based on a Wilcoxon rank-sum test with *p <* 0.05. Right: Group bootstrapped means (1000 iterations) and 95% confidence intervals for low-jar responses. Statistically significant differences between groups (*p <* 0.05, two-sided ttest with unequal variance) are noted with an asterisk. **b**. Estimated subject bias versus the maximum-likelihood estimate (MLE) of the rare-ball weight, *ρ*, for subjects best fit by the Noisy Bayesian model in asymmetric blocks. Vertical lines indicate the rare-ball weights used by ideal observer’s. **c**. Inferred subject bias versus the MLE of the response bias (Prior) for subjects best fit to the Prior Bayesian model in the asymmetric block (marker legend as in **b**). Negative values correspond to a bias in favor of the low jar. In **b** and **c**, regression lines are shown for group-blocks with significant correlations (*p <* 0.05).

These relationships were consistent with synthetic responses generated by the two model classes in parameter regimes identified by experimental data. These simulated responses also exhibited relatively high variance and low bias for the heuristic models and low variance and high bias for the suboptimal Bayesian models (Fig. S14, top). For the heuristic models, the simulated choice patterns were intrinsic to the modeled strategies and relatively insensitive to the task conditions, with similarly high variances and low biases under both the symmetric and asymmetric task conditions. For the suboptimal Bayesian models and subjects best-fit to them, the high bias reflected two forms of mistuning: 1) an under-weighting of evidence from a rare-ball observation, *ρ*, compared to the ideal observer, for the Noisy Bayesian model (Fig. 4**b**); and 2) a prior favoring the low jar for the Prior Bayesian model (Fig. 4**c**). These trends were not apparent in symmetric conditions (which also included far fewer subjects best fit by heuristic strategies; Fig. S15) and thus were consistent with the idea that suboptimal Bayesian-based inferences about asymmetric quantities yield systematic biases.

### Complexity-based ordering of bias-variance trends

To understand how these relationships between bias, variance, and strategy (Bayesian-based versus heuristic) related to the complexity of those strategies, we used a complementary, model-free analysis that measured strategic complexity in terms of the mutual information (MI) shared between each subject’s observations and their responses across trials in each block:

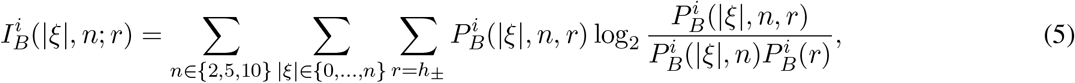

where *i* is a subject or model, *B* is the block, *n* is the size of a sample, |*ξ*| is the number of rare balls in the sample, and *r* is the corresponding vector of responses in a block (Fig. 5**a**). Following principles of rate-distortion theory [8], we compared these values to the subject’s accuracy (i.e., the fraction of correct responses within a block) and the maximum possible expected accuracy in a block with infinite trials (asymptotic bound) based on a fixed level of MI between the observations and the response (the accuracy “bound”). This approach allowed us to identify two aspects of a subject’s strategy: 1) the MI between responses and relevant observations (ball sample in the current trial) *I*; and 2) how effectively that level of MI was used to make decisions, represented by the proximity of a subject’s accuracy to the upper bound (in general high complexity/MI does not necessarily imply high accuracy, because in principle, complex strategies can use irrelevant information and/or be ineffective).

**Figure 5:**
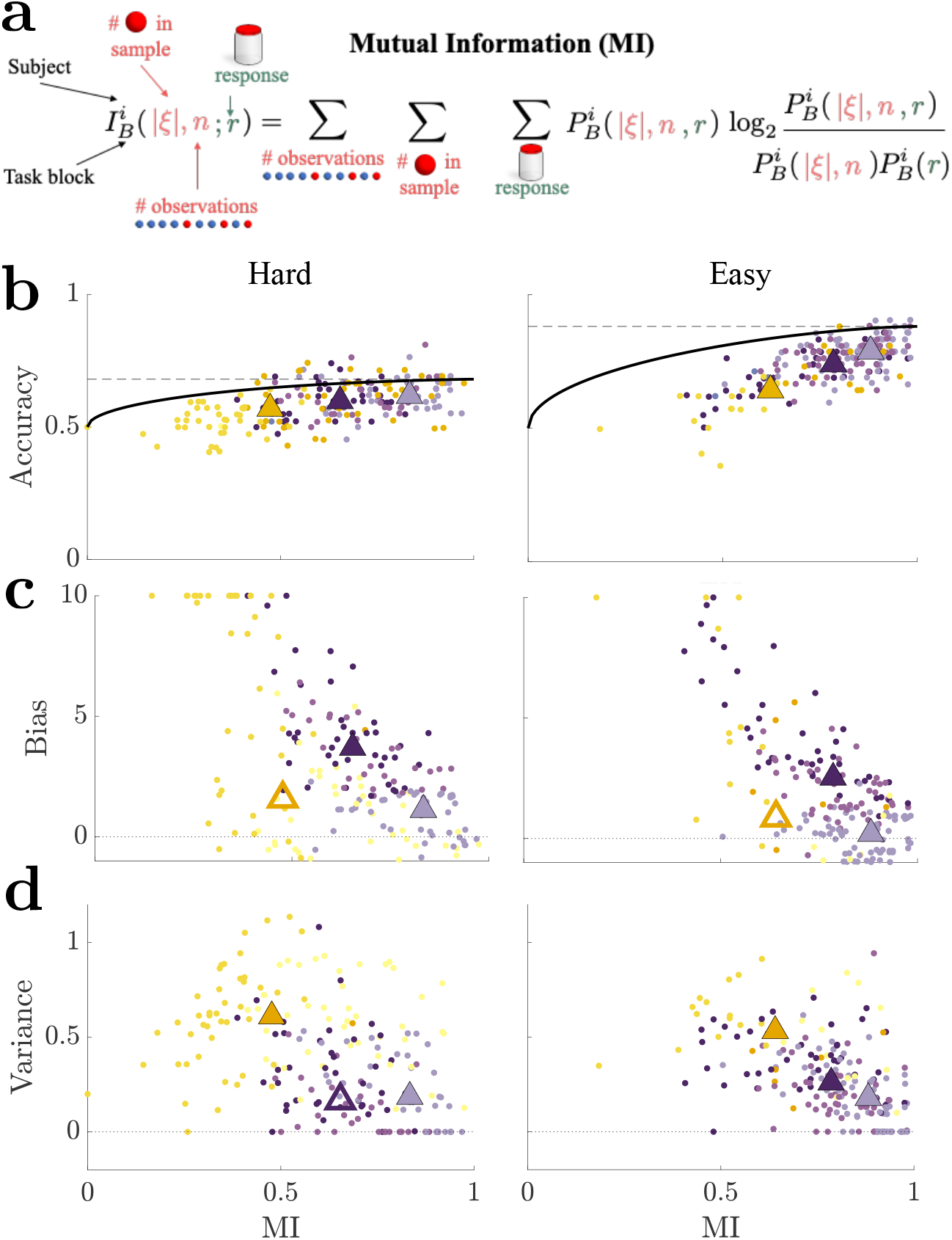
More complex but suboptimal strategies exhibit more bias. **a**. Mutual information (MI) between the number of rare balls in a sample (|*ξ*|), the sample length (*n*), and the response (*r*) for each subject and block. **b**. Accuracy versus mutual information (MI, computed as bootstrapped means from 1000 iterations per subject) for each asymmetric block (columns, as labeled). Dots represent data from individual subjects, color coded by the model that best fit a subject’s responses (colors as in Fig. 3). The bound is the maximum accuracy attainable in the limit of many trials by the optimal observer under fixed MI. Subjects with higher MI values were closer to the optimal bound. Note that points could exceed the asymptotic bound because the number of trials for each subject was finite. The dashed horizontal lines indicate the bounds on accuracies for maximum MI values. Median values for the Nearly Ideal, Suboptimal Bayesian and Heuristic groups are indicated with triangles (as in Fig. 4**a**). In each case, filled Suboptimal Bayesian and Heuristic triangles denote statistically significant differences in mutual information from the nearly ideal group (*p <* 0.05) based on a Wilcoxon rank-sum test. Median values for all 3 groups showed increase in both accuracy and MI ranking from lowest (Heuristic), middle (Suboptimal Bayesian), highest (Nearly Ideal). **c**. Relationship between bias fits and MI, triangles represented as in **b** based on statistically significant differences in bias. **d**. Relationship between variance fits and MI, triangles represented as in **b** based on statistically significant differences in variance.

In general, subjects who used more-complex strategies (in terms of their use of information from the current trial) also tended to be more accurate for all five blocks, and subjects using the most-complex strategies closely approached the optimal accuracy bound (Figs. 5**b** and S16). Moreover, both accuracy (measured in absolute terms or relative to the optimal bound) and complexity depended systematically on strategy type, such that both increased, on average, going from subjects best fit by heuristic models, to suboptimal Bayesian models (the Noisy Bayesian or Prior Bayesian model), to nearly ideal Bayesian models (the Noisy Bayesian Set *ρ* model; Fig. 5**b**). Given this increase in complexity from heuristic to Bayesian models, it followed that subjects using suboptimal strategies in the asymmetric conditions exhibited a bias-variance trade-off that was inverted relative to its typical formulation with respect to complexity: subjects using the less-complex, suboptimal heuristic strategies tended to make errors characterized by relatively high variance (and noise) but low bias, whereas subjects using the more-complex, suboptimal Bayesian-like strategies tended to make errors characterized by relatively low variance (and noise) but high bias, compared to subjects using the most-complex and near-optimal Bayesian-like strategies (Fig. 5**c** and **d**).

This complexity-based ordering of strategies, from simpler heuristics to more complex Bayesian-like strategies, was robust to two alternative metrics for complexity. The first reflected the MI between the responses and observations from the current trial, as above, but also included the subject’s choice from the previous trial as an additional “observation” to account for potential sequential choice dependencies and, more generally, the use of both relevant and irrelevant sources of information to make choices (Fig. S17). This approach tended to give a higher estimate of MI compared to the MI obtained without using the previous response for subjects best-fit by heuristic strategies, but this new MI estimate still remained lower than for Bayesian-like subjects.

For the second metric, we used a fundamentally different approach and considered algorithmic complexity [9], which has also been considered in the context of limits and trade-offs in human inference [10–12]. Unlike the MI metric, which is data-driven and requires no explicit assumptions about the underlying strategy, computing algorithmic complexity requires not only assuming a particular strategy but also assigning computational costs to each component of the strategy. To be as general as possible, we computed algorithmic complexity for each model depicted in Fig. 3 by simply counting the total number of operations (arithmetic, writing to memory, reading from memory, and storage) needed to perform the task. This metric tended to be much larger for the Bayesian compared to the heuristic strategies (Fig. S18), reaffirming our previous findings, and, while alternative reasonable costs could be assigned for each operation, these approaches would merely amplify our results.

These trends relating model complexity, bias, and variance were also apparent in simulated data using distributions of parameter values that mimicked the best subject fits (Fig. S14, middle and bottom). This result supports the idea that patterns of bias and variance are inherent to the strategies described by these models and not simply idiosyncrasies of the subjects’ behavioral patterns, with errors in more complex, Bayesian-like strategies leading to increased biases, but less-complex strategies based on the pattern of observations leading to increased variance.

## Discussion

How do people’s error tendencies depend on the inference strategies they use? We examined the properties of errors made by human subjects performing a two-alternative forced choice task with asymmetric evidence [7, 13, 14]. The evidence took the form of two colors of balls drawn from jars, such that one (“rare”) color was produced less often than the other. Similar to ideal observers, most subjects exhibited a choice asymmetry favoring the option that produced fewer rare balls. In addition, subjects fell into two categories depending on the type of strategy they used. Those using heuristic strategies, which were based on less information and fewer algorithmic operations, displayed substantially more choice variability but relatively little bias beyond the choice asymmetry of the ideal observer. In contrast, those using more-complex, suboptimal Bayesian strategies displayed minimal increases in choice variability but much more bias than the ideal observer. These effects reflected the nature of the suboptimalities introduced by each strategy type: the heuristic strategies that we considered did not attempt to capture specific task features responsible for choice asymmetries and thus tended to just add variability, whereas Bayesian-like strategies that we considered did attempt to model those features explicitly but, when implemented suboptimally by the subjects, tended to exacerbate asymmetries inherent to such decision rules.

These findings provide new insights into bias-variance trade-offs that are well established in machine learning and related fields [2, 3] and can be used to account for individual differences in human behavior under certain conditions [1, 4]. These trade-offs can be conceptualized in terms of fitting various functions that differ in complexity (e.g., polynomial order) to noisy data whose generative source is unknown. In general, simpler (e.g., linear) models tend to have higher bias because they miss higher-order (e.g., nonlinear) features of the generative source but lower variance because their best-fitting parameters are relatively stable across different data instances. In contrast, more complex (e.g., high-order polynomial) models tend to have lower bias because they can capture complex features of the data but higher variance because the specific features they capture can differ across different data instances. Critically, this traditional conceptualization is based on the assumption that each model, regardless of its complexity, is “optimal,” using the best-fitting parameters given the data and thus does not introduce additional suboptimalities and errors.

In contrast, we considered cases in which the different models (inference strategies) could differ in not just complexity but also (sub)optimality. Suboptimalities could result from the model used or a mistuning of the model parameters. The suboptimalities introduced by the two broad classes of strategies that we considered introduced errors that inverted the expected relationship between bias and variance relative to strategic complexity. One major component of this inversion was the large increase in bias for suboptimal Bayesian strategies. In general, mistuning of Bayesian model parameters is not surprising, given that Bayesian models are computationally expensive [15] and difficult to tune appropriately [6, 16]. However, the nature of this mistuning for tasks involving asymmetric evidence is different than for more commonly studied tasks involving symmetric evidence, in several ways. These differences highlight specific challenges that an effective inference strategy must overcome and can be used to predict specific patterns of errors that people are likely to make under certain conditions, as follows.

First, a major factor governing performance on inference tasks with either symmetric or asymmetric evidence is the amount and/or quality of available observations. In general, inferences based on less evidence tend to be less accurate [17, 18]. However, when the evidence and priors are symmetric, the ideal observer is not expected to show systematic biases towards a particular alternative, which is typically considered suboptimal (although such biases can arise from near-Bayesian decision strategies [19, 20]). In contrast, when evidence is limited and asymmetric, systematic choice asymmetries can be expected even for an ideal observer. As we showed, people have a very strong tendency to exacerbate these asymmetries, even when given explicit instructions that the alternatives are equally likely. Thus, systematic biases might be a general feature of inferences that must operate on limited evidence that is asymmetric, but not symmetric, in nature.

Second, effective inference requires weighting the evidence appropriately. For symmetric conditions, this weighting must be calibrated to optimize choices but in general can be effective as long as the symmetry in the evidence is maintained, even if the evidence is mis-scaled relative to the true LLR [21]. In contrast, weighting asymmetric evidence often requires much more fine tuning that, when implemented suboptimally, can give rise to systematic errors. Subjects underweighted rare evidence in our task, which may reflect a bias toward evenly weighting the evidence gleaned by each ball type, so a strong prior over even ball-weighting may pull subjects away from the ideal weight. The underweighting of rare evidence may also be related to description-experience gap theory, which distinguishes the tendency to overestimate the importance of rare events when their frequency is described and underestimate their importance when subjects learn their frequency through experience [22–25]. For our tasks, event probabilities were both described and experienced across trials, which previously has been shown to promote better decisions [26]. Nevertheless, a substantial fraction of our subjects underweighted rare-ball evidence. This deviation from description-experience gap theory could reflect differences in our task design from those used in previous studies, in that we did not include the reward component typical of previous tasks used in studies that supported that theory [23, 27, 28]. This disagreement also likely reflects the general challenge of appropriately calibrating the quantitative value of probabilistic evidence associated with rare events.

Third, many subjects used strategies that appeared to be based on subjective priors with a preference for the low jar. These findings are distinct from previous work that examined choice biases in tasks with symmetric evidence but asymmetries in expected choice frequencies [29–32] or reward outcomes [31, 33–36]. Under those conditions, biases based on asymmetric priors are common and, on average, tend to follow established, normative principles often formulated in the context of Signal Detection Theory [29] and/or sequential analysis [37]. In our study, subjects tended to either use inappropriate priors (e.g., subjects whose choices were best matched by the Prior Bayesian model with a prior biased towards the low jar) or neglect the symmetric prior altogether (e.g., subjects whose choices were best matched by heuristic models). These strategies could, in principle, reflect a relatively common form of recency bias that can cause an initial belief shift in the direction of the previous response [30, 31,33, 34, 38, 39], and, more generally, is consistent with many previous findings of mistuned priors [40–44].

The other major component of the inverted bias-variance trade-off that we identified was the relatively high variance for subjects who used heuristic versus Bayesian-like strategies. In the classic bias-variance trade-off, it is critical to distinguish variance (variability driven by sensitivity to noisy observations) from noise (variability driven by intrinsic factors): the former is expected to be anti-correlated with bias, because they represent two sides of the same underfitting/overfitting coin, whereas the latter is not. Likewise, we attempted to distinguish the two sources of choice variability in terms of: 1) psychometric slope, which we interpreted primarily as noise because it represents a general, LLR-dependent degradation of choice accuracy; and 2) the choice mean absolute error, which we interpreted primarily as variance because it represents observation-specific choice variability. Although both measures likely reflect both sources of variability, our models are consistent with our interpretation. Specifically, the Bayesian models added noise to an LLR-based decision variable, which affected the steepness of the (biased) psychometric function but less so observation-specific variability. In contrast, the heuristic models made probabilistic choices in an observation-dependent manner, which affected both the steepness of the psychometric function and the observation-specific variability. These results imply that, like for the classic bias-variance trade-off, the inverted form that we found is not just an empirical observation. Rather, it is an inherent information processing trade-off that depends on whether the suboptimal strategy operates primarily on latent (as in Bayesian-like strategies; e.g., LLR) or directly observable (as in heuristic strategies; e.g., rare ball count regardless of common ball count) properties of asymmetric environments.

By focusing on human inference on observations of asymmetrically available evidence, we uncovered a novel trend in the bias and variance properties of suboptimal inference strategies. We conjecture that this inversion of the classic bias-variance trade-off arises due to the strong tendency of people to mistune Bayesian strategies further along the direction of existing choice asymmetry. This finding also demonstrates the power of adapting Bayes-optimal models to suboptimal counterparts as a way of distinguishing strategies in a human cohort. Our study of strategy complexity also provided another way of distinguishing between Bayesian-like and heuristic models, in addition to their distinct choice error trends. In general, probing how humans make inferences in the presence of asymmetric evidence highlights relationships between bias, variance, complexity, and human error that cannot be observed in standard decision tasks and provides unique insight into the basis of human idiosyncrasies and bias-variance trade-offs for suboptimal inference strategies.

## Materials and Methods

### Experimental Design

We recruited 201 consenting subjects to perform the Jar-Discrimination Task on the Amazon Mechanical Turk crowd-sourcing platform (95 female, 105 male, 1 non-disclosed). Subjects were recruited only if they had a 95% or better approval rating and had performed at least 100 previous approved tasks and were compensated $4.50 for completing the task. Subject location was restricted to the United States. Human subject protocols were approved by the University of Pennsylvania Internal Review Board. The task and some of the analyses were preregistered at osf.io prior to data acquisition (doi:10.17605/OSF.IO/J9XET). Analyses presented in Fig. 2**b,c**, Fig. S3, Fig. S4, Fig. S12 and Fig. 3 and the MLEs from Fig. 4**b,c** were performed exactly as listed in the preregistration.

#### Task Design

The goal of the task was to identify which of two jars was the source of a sample of balls shown to the observer. The jars were equally likely to be the source *a priori*. On each trial, subjects were shown a sample of 2, 5, or 10 red and/or blue balls drawn randomly with replacement and asked to determine which of the two jars displayed on the screen was the source of the sample (Fig. 1**a** and Fig. S1). Subjects were informed that the two jars were *a priori* equally likely to be the source. The ratios of colored balls in each jar were varied to create five blocks of trials and could be described by the proportion of balls of one color, termed the “rare-ball” color. The rare-ball color remained consistent throughout all blocks. Blocks were defined by the following rare-ball fractions for the high jar (containing more rare balls)/low jar (containing fewer rare balls): Control (0.9/0.1), Hard Asymmetric (HA; 0.2/0.1), Hard Symmetric (HS; 0.55/0.45), Easy Asymmetric (EA; 0.4/0.1), Easy Symmetric (ES; 0.7/0.3).

Before beginning the full task, subjects were shown a training slideshow and performed 24 trials in the control block. To continue to the full task, each subject was required to respond correctly on at least 80% of the control trials. Full sessions included randomized block orders for the remaining 4 test blocks interspersed with 12 control trials between test blocks. Each test block consisted of 42 trials, with randomly ordered but equally sampled values of: 1) the jar used for ball draws, and 2) sample length for each trial (2, 5, or 10 balls).

Prior to data acquisition, we used synthetic data generated by simulating the responses from the proposed models to confirm that models were identifiable given the task conditions and amount of data to be collected (Fig. S9). We determined the number of trials in a block by balancing: 1) model parameter identifiability, with 2) reasonable task-time length for human subjects (i.e., about 30 min per session). The jar ratios were selected based on generated synthetic responses of the ideal observer, such that overall accuracy was matched between the asymmetric and symmetric blocks at each difficulty (i.e., the hard asymmetric and hard symmetric tasks were matched in accuracy). Models were developed and fit to pilot data to ensure model and parameter identifiability (See Model Fitting and Comparison below).

### Psychometric Functions

Subject response data for each block were fit by a three-parameter logistic function:

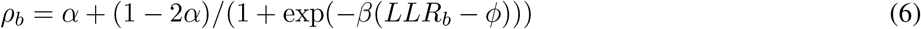

where *α* is the lapse rate, *φ* is the LLR value at which each choice (high or low jar) is equally likely, and *β* is the slope around the point *φ*. Bias was defined as a non-zero value of *φ*, so that positive (negative) values correspond to biases towards (away) from the low jar. Noise was defined as 1/|*β*|, so that shallower functions correspond to higher noise. Variance was defined as the weighted average of the absolute value of the residuals (mean absolute error),

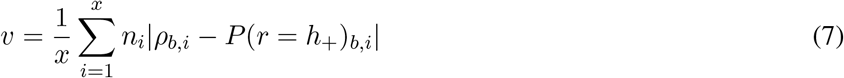

where *x* is the number of LLR values for a block, *n*_*i*_ is the number of trials at a given LLR value, *ρ*_*b,i*_ is the logistic fit for a given block-LLR, and *P*(*r* = *h*_+_)_*b,i*_ is the probability of a high jar response from the observer for a given block-LLR. Larger values of *v* reflected more variance.

Our interpretation is based on the idea that noise is driven by errors in the internal representation of the LLR, while variance reflects an observation-specific, rather than LLR-based, strategy. Based on the two model classes studied here (see Models below), we find that models that rely on the LLR (Bayesian models) and the subjects best fit to them are fit with some noise but substantially less variance compared to models and subjects that use a pattern-based approach (heuristic models, Fig. S19). This latter group also shows substantially larger values for noise, which reflect the the poor logistic fits to these responses (e.g., Fig. S6 grey traces).

### Models

#### Bayesian Models

One class of models we considered depended on the probabilities of ball samples coming from the high or low jar that would be computed by a Bayesian observer.

##### Ideal Observer

In the simplest case without noise, an ideal Bayesian observer makes a decision based on a sample of *n* balls drawn from one of the jars, *ξ*_1:*n*_, where *ξ*_*i*_ = 1 (*ξ*_*i*_ = *−*1) denote an observation of a rare (common) ball color. The ideal observer uses these observations to update the log likelihood ratio (*belief*), 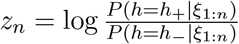, between the probabilities that the sample came from a jar with a rare ball frequency of *h* = *h*_+_ (high) or *h* = *h*_*−*_ (low). We can write the update iteratively:

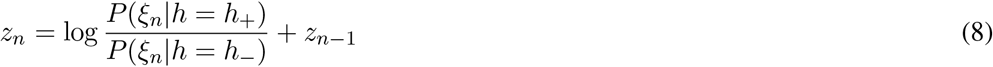

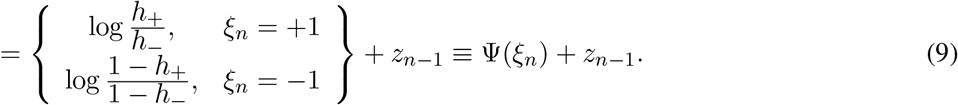

The most likely choice based on *n* ball draws is given by the sign of *z*_*n*_ (*z*_*n*_ *>* 0 → choose the high jar; *z*_*n*_ *<* 0 → choose the low jar). In all blocks, the probability that either jar was the source of the sample was 0.5, so that the ideal observer model had a flat prior, and 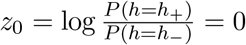.

In symmetric environments, *h*_+_ = 1 *− h*_*−*_, so

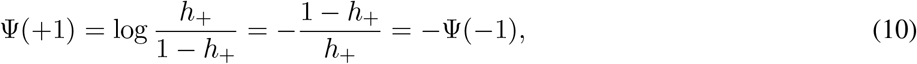

and thus the magnitude of the belief increment is the same for either observation (|Ψ(+1)| = |Ψ(*−*1)|). When the environment is asymmetric, *h*_−_ *<* 1 − *h*_+_, and different ball colors correspond to different evidence weights (|Ψ(+1)| *≠* |Ψ(*−*1)|).

For *n* ball draws, we can compute the probability of the responses (choices) on a given trial, *r* = *h*_*−*_ and *r* = *h*_+_ for the low and high fraction jars as

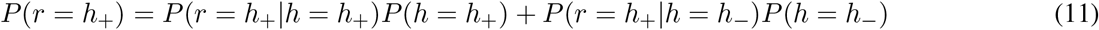

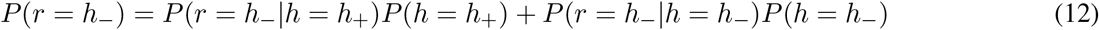

using binomial distributions. For example,

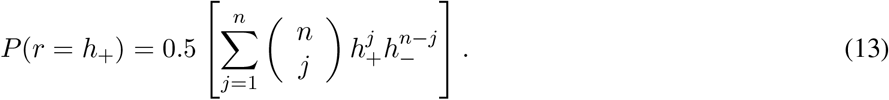

##### Noisy Bayesian

We extended the ideal observer model to include noisy belief updates, with means and variances of arbitrary magnitude. To do so we let *w*_*n*_ ∼ 𝒩 (0, *a*^2^) be a normally distributed random variable with zero mean and variance *a*^2^ that was fit as a free parameter. We defined the weight updates by

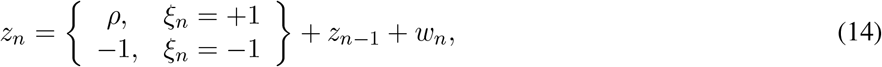

where *ρ* is a free parameter representing the belief update in response to observing a rare ball, *ξ*_*n*_ = 1. Because the sign of *z*_*n*_ is all that matters for determining a model observer’s response, we normalized the update in response to a common ball to remove an unnecessary parameter. Thus, fits using this model had two free parameters: *a*^2^ and *ρ*.

##### Noisy Bayesian Model Set ρ

For this model, the belief updates are given by

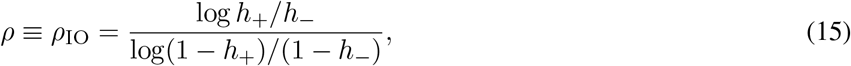

and equal those in a rescaled ideal Bayesian model. Each belief update is perturbed additively by a Gaussian random variable with variance, *a*^2^. We set *ρ* to the optimal value *ρ*_IO_, and thus the variance, *a*^2^, was the only free parameter.

##### Prior Bayesian Model

We modified the Noisy Bayesian Model Set *ρ* to include a free parameter *z*_0_ for the prior. An observer using this model uses potentially unequal prior probabilities,

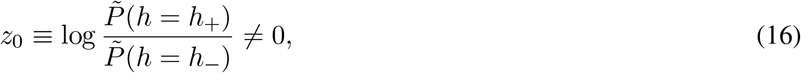

where 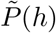 represents the observer’s assumed prior probability, which may differ from the the true prior probability that a jar with rare ball fraction *h* is a source of the sample. A positive (negative) value of *z*_0_ implies that the observer believes *a priori* that the high (low) jar is more likely to be the source of a sample. Thus, fits using this model had two free parameters: *a*^2^ and *z*_0_.

#### Heuristic Models

The other class of models (heuristic) did not depend on the likelihood functions associated with drawing a ball of a certain color from either jar.

##### Variable Rare Ball Model

The probability of choosing either jar in the Variable Rare Ball Model depends only on whether a certain number rare balls (*θ*) are observed in a sample in the current trial (*N*),

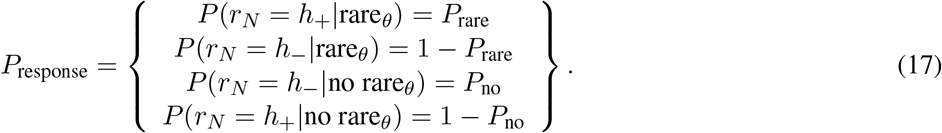

Here *r*_*N*_ is the response on the current trial *N*, 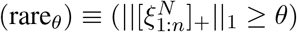 corresponds to observing *θ* or more rare balls (or the sum of positive entries of 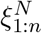 being at least *θ*), and 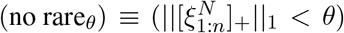 to observing no rare balls in the current trial (or the sum of positive entries of 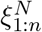 being less than *θ*). Thus, fits using this model had three free parameters: *θ, P*_rare_, and *P*_no_.

##### Rare Ball

For this model we assumed that *θ* = 1, reducing the number of free parameters to two.

##### Guess Model

In this model, the probability of each choice is fixed, and independent of the sample. The Guess Model includes one free parameter that determines the probability of choosing the high jar:

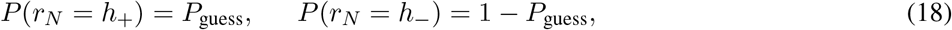

regardless of any observations within a trial.

##### Alternative (unused) models

In addition to the above models, we considered four alternative models, three Bayesian and one heuristic. The Bayesian models included a variation of the Noisy Bayesian with a bias in the prior probability of the two choices (3 free parameters) and a history-dependent model with asymmetry in favor of low jar responses, but we found neither of these to be identifiable (see Model Fitting and Comparison below and Fig. S10). We also considered a windowing Bayesian model, in which a specified amount of evidence was used consistently across trials (with the observer drawing from previous trials if the evidence on the current trial was insufficient), and a history-dependent rare ball model, in which the probability of a choice depends on observing a rare ball in the sample, *and* the choice *r*_*N−*1_ on the previous trial. In both cases, fewer than 5 subjects per block were fit to these models (Window: CT-1, HA-1, HS-3, EA-2, ES-3; Hist.-Dep: HA-1, HS-1) and were not included in further analyses. Subjects originally fit to these models were refit with accepted models listed above with history-dependent subjects fitting to guess models and windowing subjects fitting to a variety of Bayesian and heuristic strategies (7 Bayesian, 3 heuristic fits)

### Model Fitting and Comparison

#### Parameter fitting

We fit model parameters to data using Bayesian maximum-likelihood estimation. We obtained the posteriors over the parameters by considering the vectors of responses, *r*_1:42_, and observation samples, ***ξ***_1:42_, across all 42 trials in a block (***ξ***_1:60_ for the control block that had 60 trials). For instance, to infer the noise variance, *a*^2^, and rare ball weight, *ρ*, in the Noisy Bayesian Model, we applied Bayes’ rule and then computed the probability of a response *r*_*N*_ in a given trial conditioned on observations ***ξ***_*N*_ as

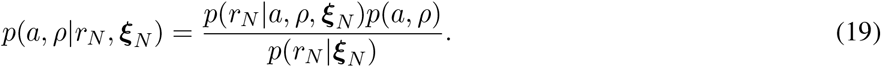

Because the denominator provides only a normalization of the probability densities of *a* and *ρ*, the primary contributions are the probability of a response *r*_*N*_ given the parameters and observations, and the prior over the parameters, *p*(*a, ρ*). We explain the choice of priors below. Because all models were defined in terms of either simple binary random variables or thresholded Gaussians, we could evaluate the associated likelihood functions analytically. For instance, in the case of the Noisy Bayesian model, for a trial with 5 balls, and a sample containing 4 common and 1 rare ball (***ξ***_*N*_ = (*−*1, *−*1, +1, *−*1, *−*1)), the probability of choosing the high jar, *r*_*N*_ = *H*, is

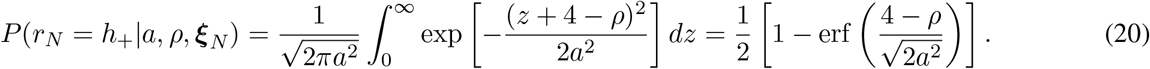

For models in which responses are independent across trials, we used the trial-wise response probabilities to compute the posteriors given responses and samples in a block of trials,

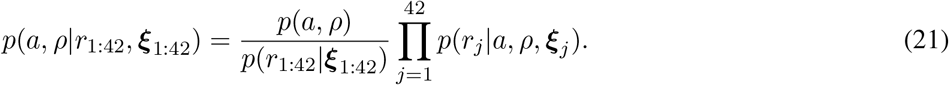

The maximum of this posterior is the maximum likelihood estimate of the model parameters. The interval of parameters containing at least 95% of the maximum likelihood estimate were included as credible intervals for the model fits.

#### Determining model identifiability

To design the human task and determine whether the models are identifiable from the given data, we performed model comparisons on synthetic data. We first used the Noisy Bayesian model to determine the minimum number of trials (42) needed to fit synthetic data and produce a task with a reasonable task duration for online data acquisition (30 minutes or less) (Fig. S9). However, for this model, parameters produced with a flat prior were not always identifiable, given the amount of data that we could reasonably expect to collect in a block. This problem resulted from dependencies between the noise variance, *a*^2^, and rare ball weight, *ρ*, parameters (Fig. S9 HA). To account for this effect, we used pilot data from 20 subjects to create an informative prior based on the subjects’ posteriors. The informative priors were computed as a smoothed version of averaged posteriors produced by the pilot subject’s fits to the Noisy Bayesian model. The averaged posterior was smoothed with respect to each parameter (Fig. S20). To weaken the posterior with respect to *ρ*, the averaged marginal posterior was filtered using a normal distribution 𝒩 (*µ, σ*^2^) where the mean *µ* was set at the maximum value of the averaged marginal posterior and the variance *σ*^2^ was set such that the median mean squared error (MSE) of the parameter fits *ρ* for 100 synthetic Noisy Bayesian datasets was below one. The averaged marginal posterior with respect to the noise parameter *a* was smoothed using the function (*x* + *c*)/(1 + *cL*) where *x* is the marginal posterior and *c* and *L* are scaling constants selected such that the averaged posterior was smooth (no jagged edges) but did not impact the accuracy of the rare ball parameter fitting (values provided in Fig. S9). Given that a low-noise parameter *a* was identified for most pilot subjects and that higher values of *a* could correspond with underweighting the value of *ρ*, we prioritized accurately identifying *ρ*.

To confirm that the new informative priors produced realistic fits for all of our Bayesian models, we applied the informative priors to model fits for data of 100 synthetically generated subjects with randomly selected model parameters for each model per block and found that the credible intervals usually contained the true parameters (median MSE *ρ* range 0.01–0.82, median MSE *a* range 0.006–0.69) (Fig. S21). The fits of the synthetic subjects also matched the averaged posteriors from the pilot data, with a strong preference for low-noise parameter values and values at or below the true rare-ball weight. Thus, using informative priors did limit identifiability at high noise variance and rare-ball weight values, but we ensured that our models could be correctly identified near or below the values predicted by the ideal observer, as was suggested by the pilot data.

We then generated model responses from 100 randomly sampled versions for each of the candidate models (sampling from the informative priors) to confirm that each model could be appropriately selected when compared to other models. We performed model comparison and selection using log Bayes factors, comparing the likelihood that a particular dataset came from one of two models by computing the log likelihood ratio of the marginal likelihoods for any given pair of models,

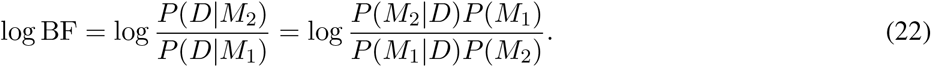

Here *D* is the data from a block of trials (*r*_1:42_ and ***ξ***_1:42_), and *M*_1_ and *M*_2_ are two models from the list we described above. For example, to compare the Noisy Bayesian Model to the Prior Bayesian Model for a given block, we integrate the probability of responses conditioned on observations and parameters against the priors over the model parameters:

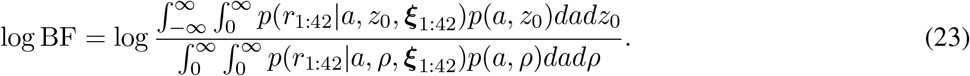

For all comparisons, we used the Noisy Bayesian model as the baseline model (in the denominator of the Bayes factor); i.e., model 1 (*M*_1_). We found that two candidate models were not identifiable as listed above (one assumed an asymmetric repetition bias, the other included a biased prior and free parameter for rare ball weight; Bayes factors correctly selected the true model *<* 80% of the time) and thus were excluded from our analyses.

#### Subject model selection

To determine the model that was most consistent with a human subject’s responses on a particular block, we computed the log Bayes factors between each alternative model and the Noisy Bayesian Model. Positive values of the log Bayes factor provided evidence in favor of a particular alternative model over the noisy Bayesian model, with evidence growing with the magnitude of the factor (we chose |log BF| *>* 1 to indicate sufficiently strong evidence in favor of a model [45]). The most-likely model was selected based on the maximal log Bayes factor value across all alternative models. If no values were *>*0, the Noisy Bayesian model was selected.

### Rate-Distortion Theory

We applied rate-distortion theory to compare the subjects’ accuracy (fraction correct) to the maximal accuracy bound obtainable by an ideal observer constrained to a fixed amount of mutual information (MI) between an observer’s response, *r* and the observation on a trial. We describe this observation as a random variable (|*ξ*|, *n*), where *n* is the size of a sample, and |*ξ*| is the number of rare balls in the sample, as:

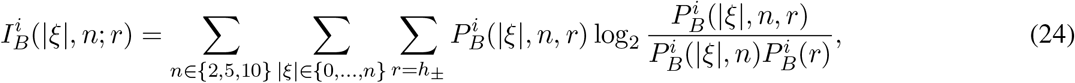

where *i* is a subject or model, *B* is the block. We computed the probabilities 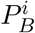 empirically.

To obtain subject estimates we used all response and observation data for the 42 trials within a block, so any particular observation sample not seen was not included in the sum. Each subject’s trials within a block were bootstrapped by uniformly resampling the data 1000 times to obtain a distribution of MI and accuracy estimates for the block.

The MI with the inclusion of the previous trial was defined as:

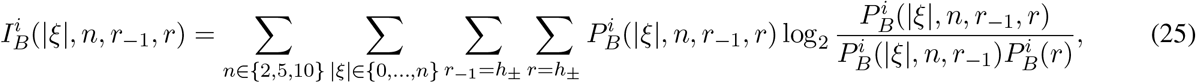

where *r*_*−*1_ is the previous trial’s response.

To define the accuracy bound for an optimal observer, we computed MI in the limit of many samples (not just 42 per block), allowing for a calculation directly using probability mass functions. As such, we considered all possible samples 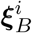, disregarding ball order, (*n* + 1 possible counts for trials with *n* = 2, 5, 10 ball draws) in Ξ and responses 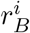 in *R* :

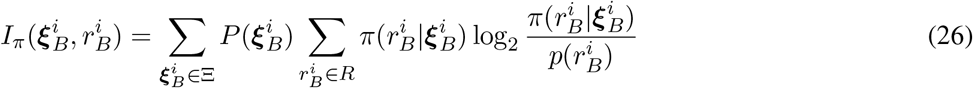

where 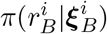 is the policy used to generate responses from observations across the block. Note, that this is simply given by the standard ideal observer model defined above when fixing the MI to unity. However, for values of MI less than one, we employed an optimization procedure, which we describe below, in order to obtain the optimal policy that uses a fixed MI budget.

#### Computing the Optimal Bound

The rate-distortion bound can be computed according to a constrained-optimization problem in which we identify the maximum possible accuracy for a given level of MI in the limit of many trials. In the ideal observer case, the policy applied to compute MI and accuracy is:

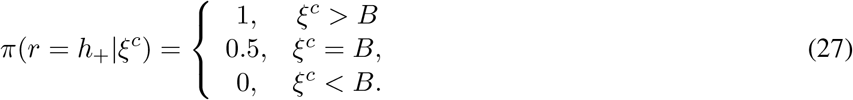

(and *π*(*r* = *h*_*−*_ |*ξ*^*c*^) = 1 *− π*(*r* = *h*_+_ |*ξ*^*c*^)) where *ξ*^*c*^ ∈ {0, 1, 2, …, *n*} is the count of rare balls observed and *B* is the number of rare balls required to trigger a high jar response. Note this provides a specific accuracy bound for a fixed value of MI, corresponding to the ideal observer. Additionally, we must compute the predictive accuracy using the value function applied to a particular policy *π*

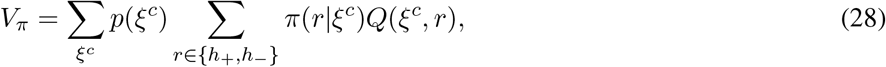

which sums over all possible combinations of unordered sample counts (*ξ*^*c*^ = 0, 1, 2, …, *n* rare balls for *n* = 2, 5, 10 balls in a trial) for which we can always compute the trial specific value function from the ideal observer 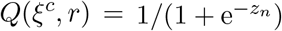, where *z*_*n*_ is the ideal observer’s log likelihood ratio.

Thus, to bound accuracy for a given MI (*I*_*π*_ ≡ *C*), we maximized the value function according to the best policy that uses the prescribed MI:

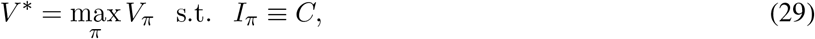

which generates the optimal predicted bounds. This maximization problem was solved using MATLAB’s constrained optimization package (`fmincon`) with a constraint given by *I*_*π*_ *≡ C* and *V*_*π*_ as the objective function.

### Algorithmic Complexity

As in [9], algorithmic complexity is described by the number of operations requires for each strategy, broken into 4 types: 1) arithmetic (*A*), 2) written into memory (*W*), 3) stored in memory (*S*), 4) read from memory (*R*). Thus complexity is defined as

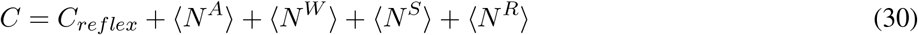

where *C*_*reflex*_ is the reflexive cost, constant across models. ⟨N^*i*^⟩ are the 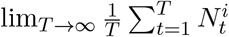 for each operator type. Details on the operations counted for each strategy are found in Fig. S18.

### Statistics

Population statistics were computed by uniformly bootstrapping 1000 times from each data set, using the same number of samples as the original sample, to identify the mean and confidence intervals. Subject p-value statistics were computed using bootstrapping and then applying the Benjamini-Hochberg correction for multiple comparisons [46]. Differences between medians were computed using a two-sided Wilcoxon rank-sum test. We defined significance as *p <* 0.05.

## Acknowledgements

We would like to thank Ronald van den Berg for his valuable feedback on the manuscript.

## Author Contributions

Conceptualization: TLE, JIG, KJ, ZPK

Methodology: TLE, JIG, KJ, ZPK

Data Collection: JIG

Investigation: TLE

Visualization: TLE

Supervision: JIG, KJ, ZPK

Writing-original draft: TLE

Writing-review and editing: TLE, JIG, KJ, ZPK

## Supplementary Materials

**Figure S1:**
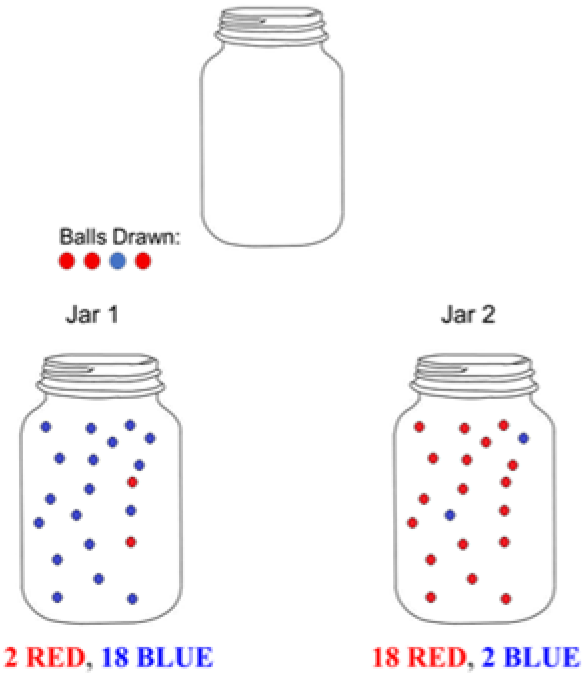
Example of the screen viewed by subjects on Amazon Mechanical Turk. A prompt at the bottom of the screen indicated to the subject to select the jar from which the sample was drawn.

**Figure S2:**
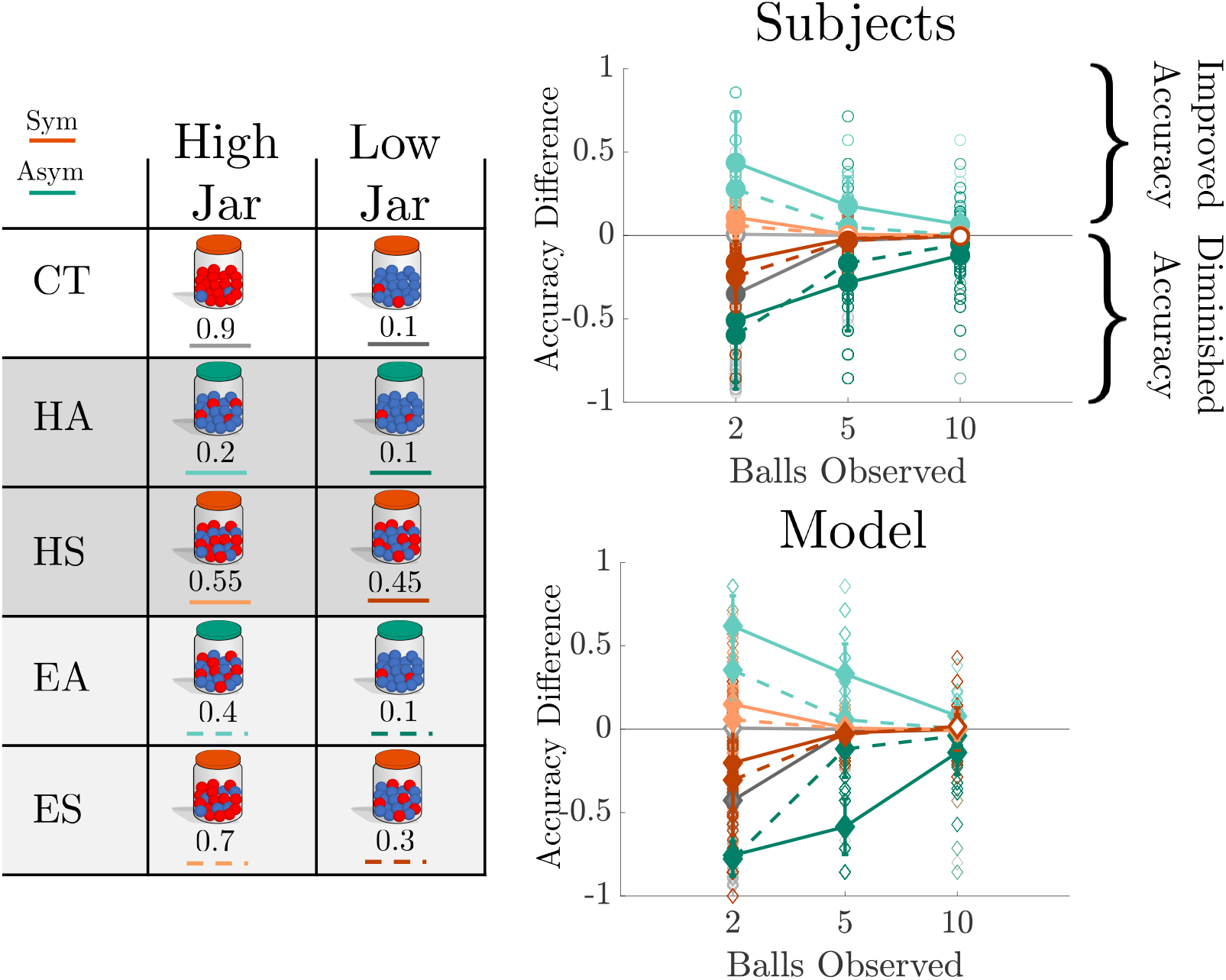
Difference in performance on trials with at least one rare ball compared to performance from all trials for the given condition for subject (top) and ideal observer (bottom) responses, plotted as a function of the number of balls observed (trial length) for each condition. Differences in accuracy that corresponded to response asymmetries tended to be larger on asymmetric blocks with fewer observations for both human and ideal observers. Individual circles (diamonds) represent individual subjects (sample-matched ideal-observer models). Summary statistics (connected by lines) were based on bootstrapped mean and 95% confidence intervals. Filled markers denote a significant population shift away from 0 (*p* < 0.05).

**Figure S3:**
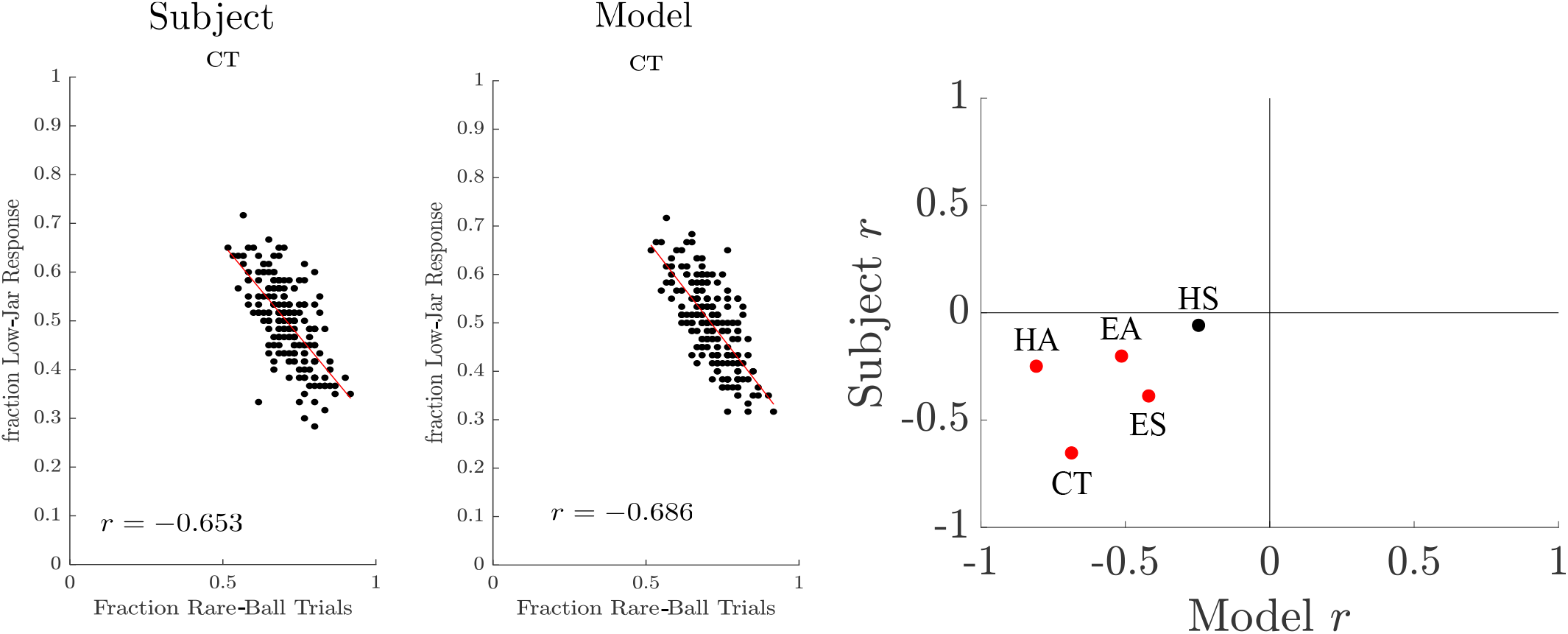
Left: Example session showing that the fraction of rare-ball trials was negatively correlated with low-jar response frequency for individual subjects and matched ideal observers (black dots). Red lines are linear fits; non-parametric correlation coefficients (*r*) are also shown. Right: Correlation coefficients *r* for the model and subjects in each block. Blocks in which both the subjects and model had significant correlations (*p* < 0.05) are noted in red.

**Figure S4:**
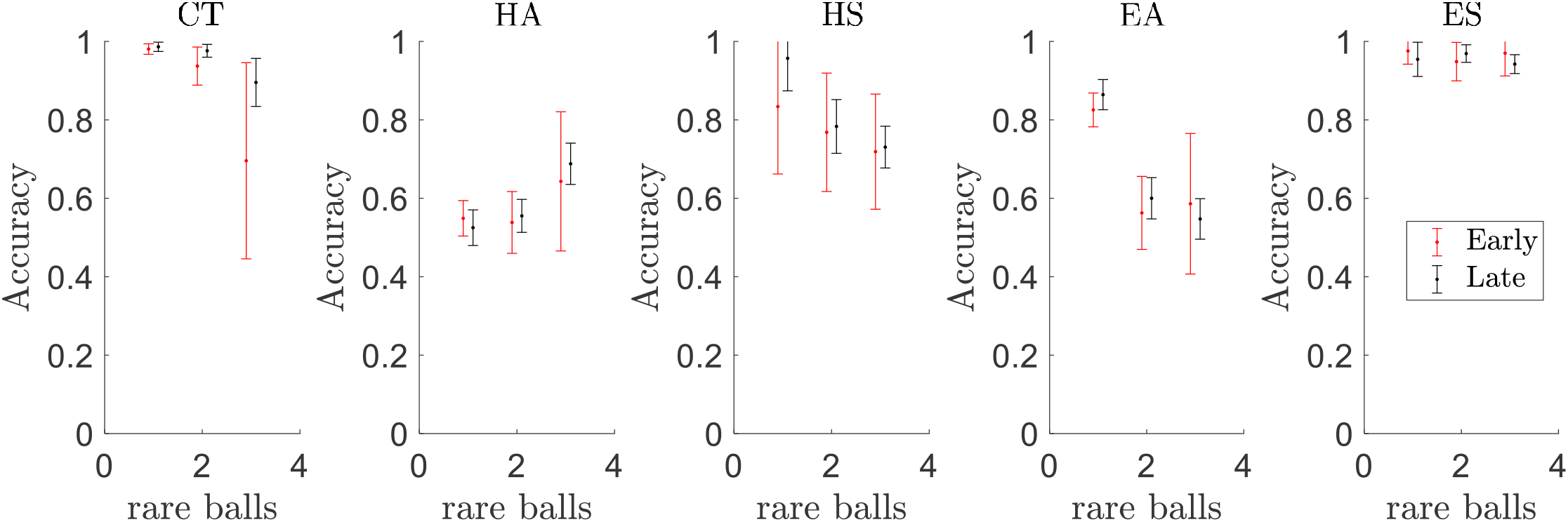
Fraction correct for 10-ball trials with 1, 2, or 3 rare balls, where the final rare ball occurred early (in the first 5 balls, observing from left to right) or late (in the last 5 balls) in the sequence of balls shown on the current trial. We found no systematic difference between the early and late trials, as evidenced by the bootstrapped population means and 95% confidence intervals.

**Figure S5:**
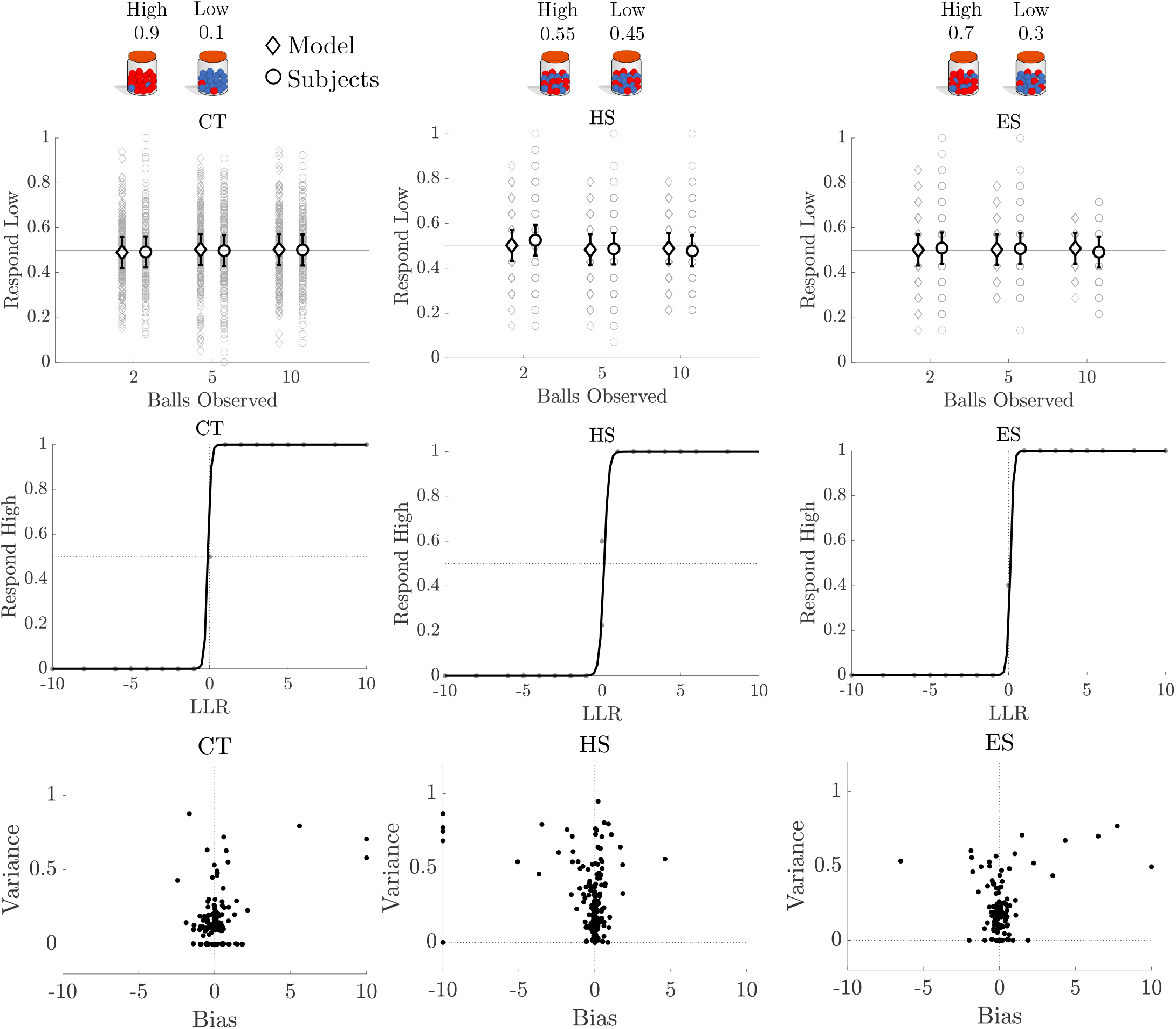
Top: Low-jar response fractions for the symmetric blocks for ideal observers (model) and subjects. Bold points and errobars are bootstrapped means and 95% CIs. Middle: Median subject responses (points) and best-fitting logistic regression functions, which were steep and unbiased as for the ideal observer. Bottom: Bias (psychometric shift) and variance (mean absolute error) from logistic fits to data from individual subjects (points).

**Figure S6:**
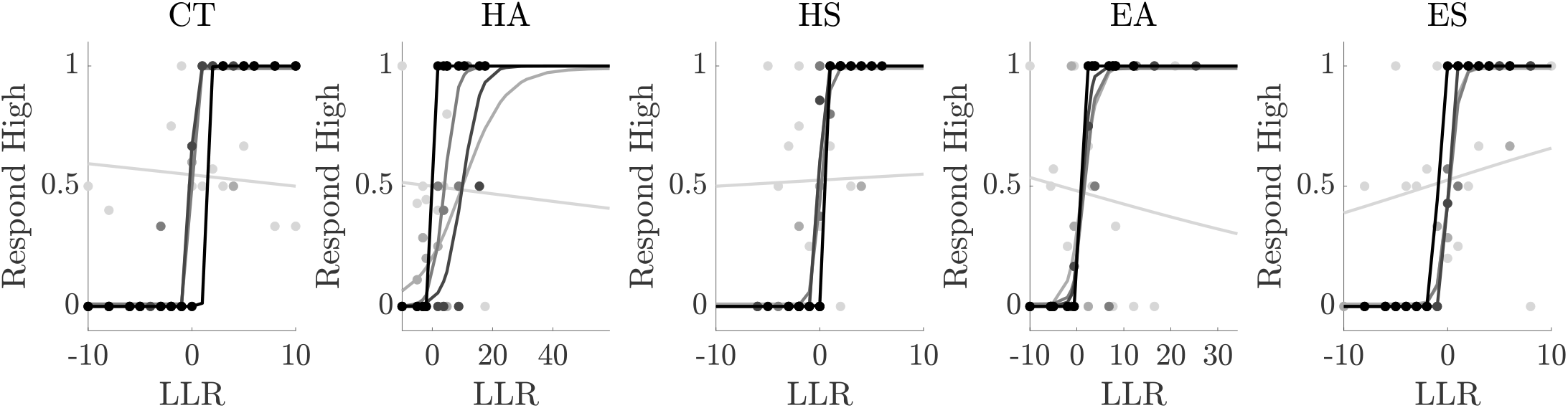
Example psychometric functions for individual subjects. Subjects were selected by the quality of their fit (identified by the log likelihood), here ranging from best fit (black) to worst fit (light grey), with intermediate fits at 25, 50, and 75th percentile rankings.

**Figure S7:**
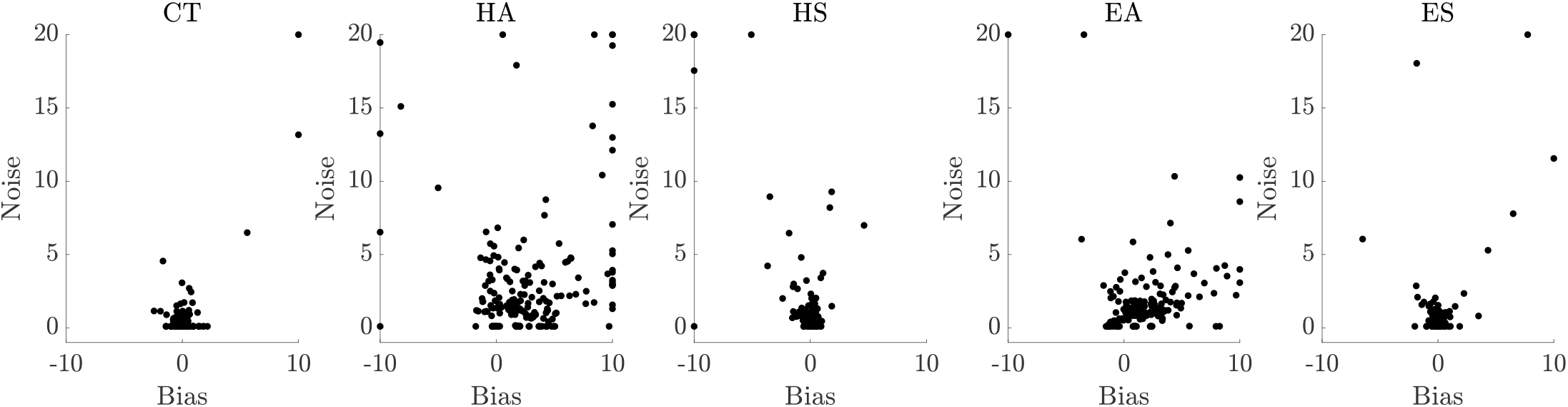
Bias-noise plots for each subject (points) and block (column). Bias and noise were determined from the offset and slope, respectively, of best-fitting logistic psychometric functions. Large noise values (> 20) were rescaled to 20 for visualization purposes.

**Figure S8:**
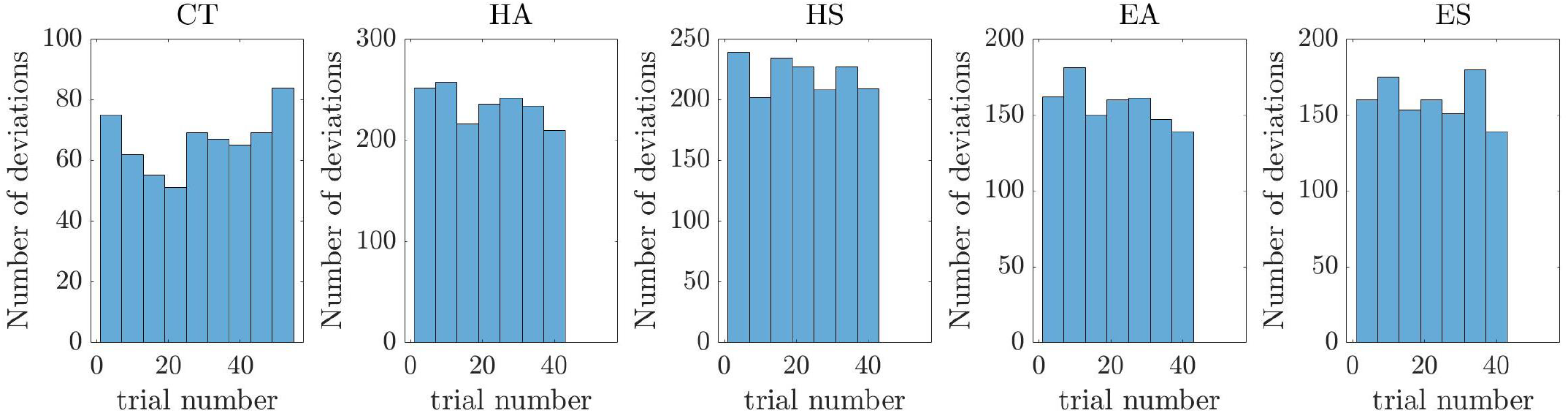
Histogram of the frequency with which subject responses deviated from the ideal observer based on trial number within a block, binned for every 6 trials. A *t*–test of deviations in the first half versus the second half of the block using bootstrapped data showed no significant difference for any block.

**Figure S9:**
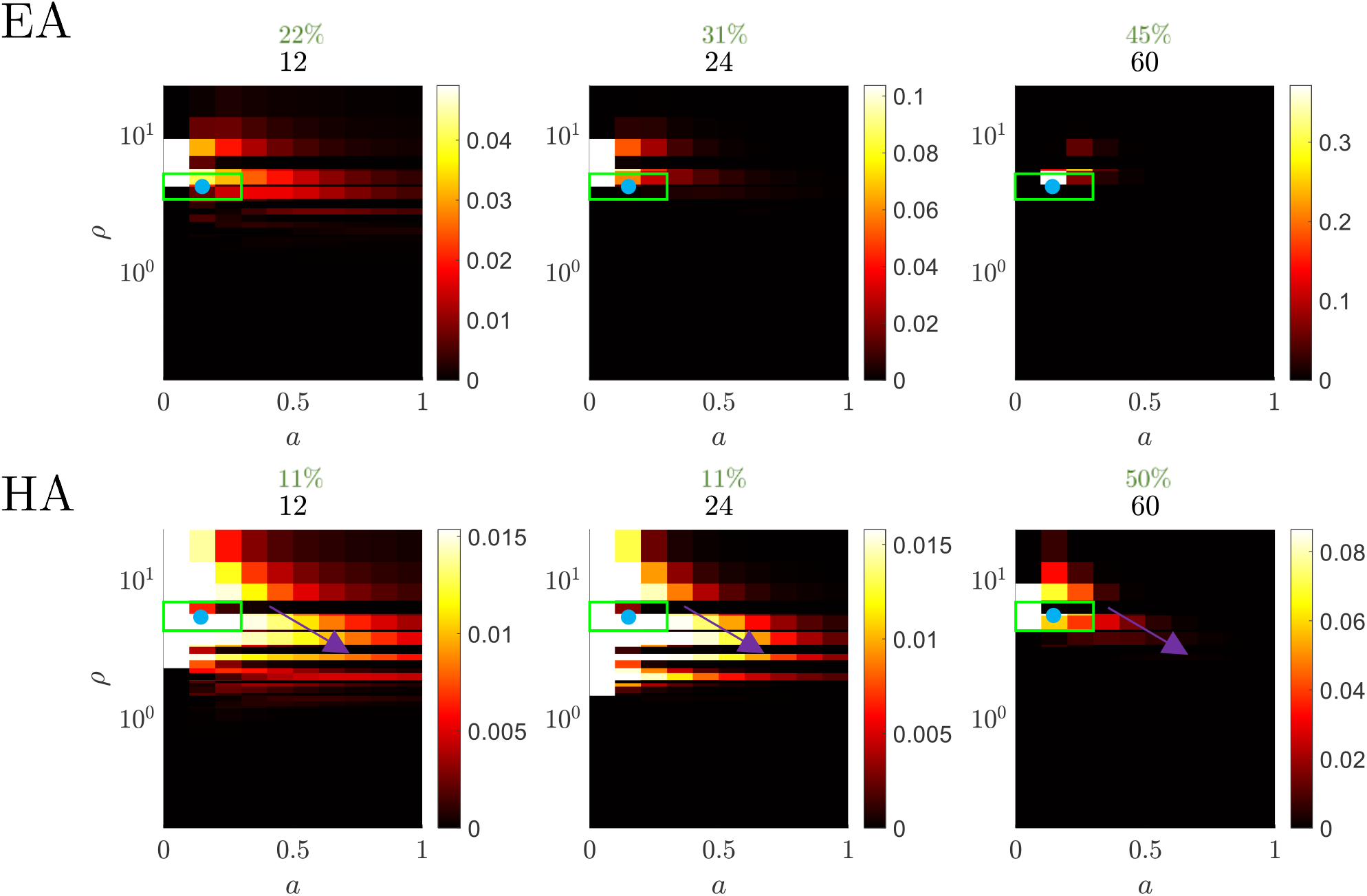
Example Bayesian parametric posteriors for the rare-ball weight *ρ* and noise variance *a* of the noisy Bayesian model using Bayesian sequential updating based on a flat prior over 0 ≤ *a* ≤ 1 and 0 < *ρ* ≤ 24.16 (computed from 0.01 ≤ *h*_*±*_ ≤ 1). Fits are based on synthetic data collected from varied block lengths (12, 24, and 60 trials, columns) of the asymmetric blocks based on responses produced using the ideal observer’s *ρ* and a low level of noise (*a* = 0.1). True parameters are shown as blue dots. By 60 trials, the parameters are well identified, with over 40% of the posterior falling within a range of one parameter value (green box, corresponding percentages shown in green). Because a flat prior is used, there can be a high likelihood for alternative scenarios in which there is a trade-off between higher noise and lower *ρ* values, as shown by the arrows in the HA fits.

**Figure S10:**
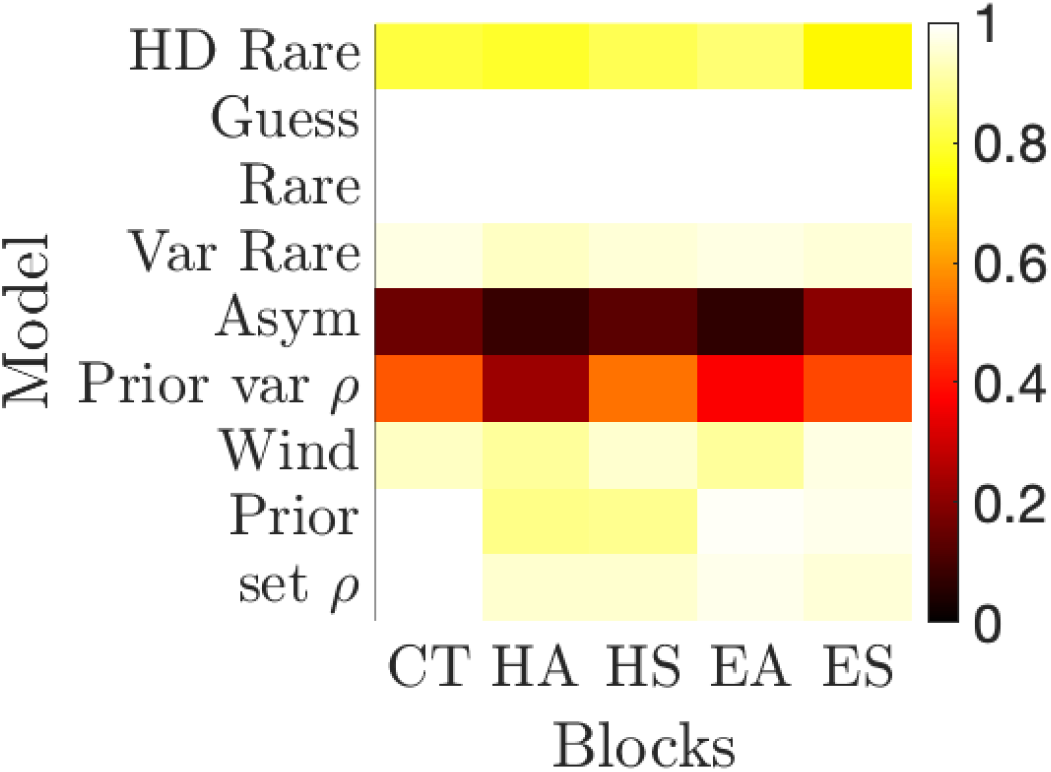
Fraction of times an alternative model was correctly identified as compared to the Noisy Bayesian model. Data from 100 synthetic subjects were produced using the human task structure (4 blocks with 42 trials, control block with 60 trials). Models are labeled and defined as in Fig. 3, except for two models that were not identifiable above 80% for all blocks (“Asym”, which assumed an asymmetric repetition bias following a low-jar response, and “Prior var *ρ*”, which was the noisy Bayesian model with biased prior) and two models that were matched to less than 5 subjects per block (“Wind”, which assumed a set window of evidence for each trial, and “HD rare”, which includes the past trial’s response in the rare ball model). These models were not used in further analyses. See the Models section of Methods for a detailed description of all other models.

**Figure S11:**
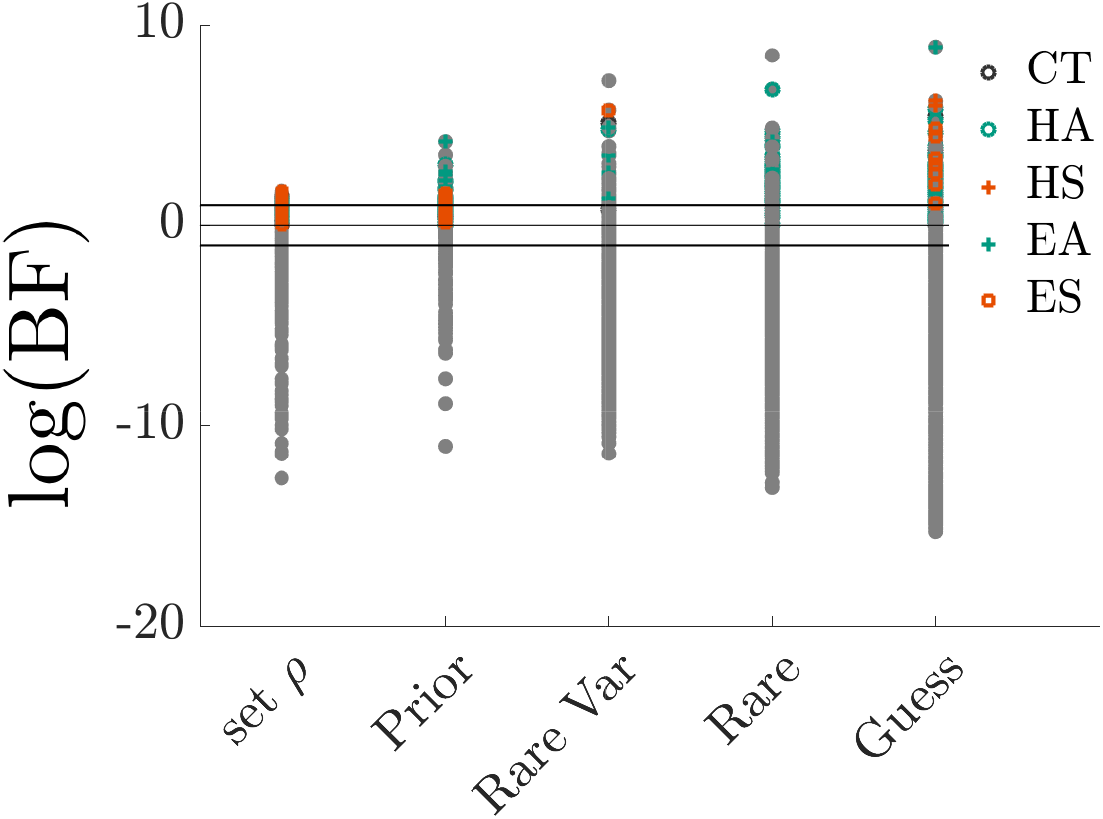
Log Bayes factors for each subject-block, computed between each alternative model and the Noisy Bayesian model. log(BF)> 0 indicates the alternative model is more likely, with log(BF)> 1 or < −1 (black lines) considered to be strong evidence in favor of the preferred model [45]. Colored (grey) dots indicate the associated model is (is not) the most likely model.

**Figure S12:**
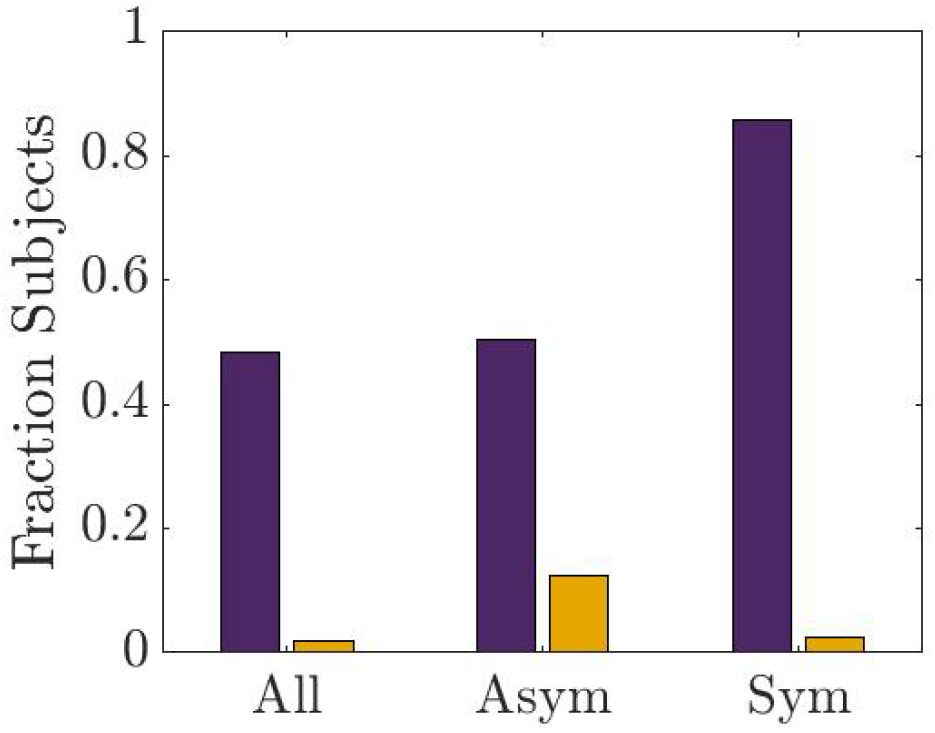
Fraction of subjects who were well matched to models in the same class, Bayesian (purple) or heuristic (yellow), across all blocks (All), asymmetric blocks (Asym) or symmetric blocks (Sym). Bootstrapped fractions of Bayesian and heuristic models are all significanty different from one another (*p* < 0.05).

**Figure S13:**
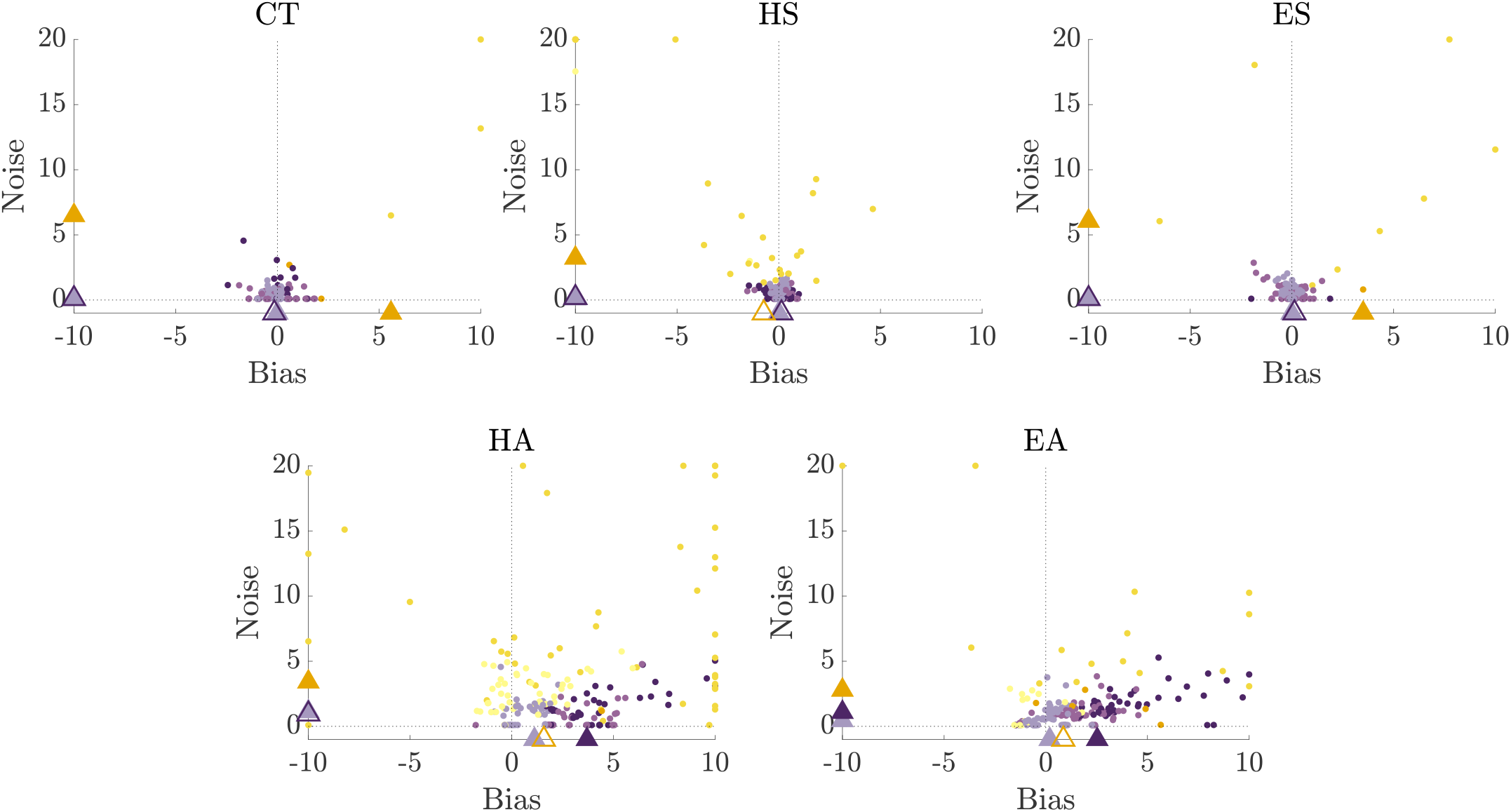
Bias-noise plots for task blocks, color coded by each subject’s best-fit model as in Fig. 3. Triangles represent medians for the the nearly ideal subjects (light purple), suboptimal Bayesian subjects (dark purple), and heuristic subjects (yellow). Filled triangles significantly differ from the nearly ideal subjects based on a two-sided Wilcoxon rank-sum test with *p* < 0.05. Large noise values (> 20) were rescaled to 20 for visualization purposes.

**Figure S14:**
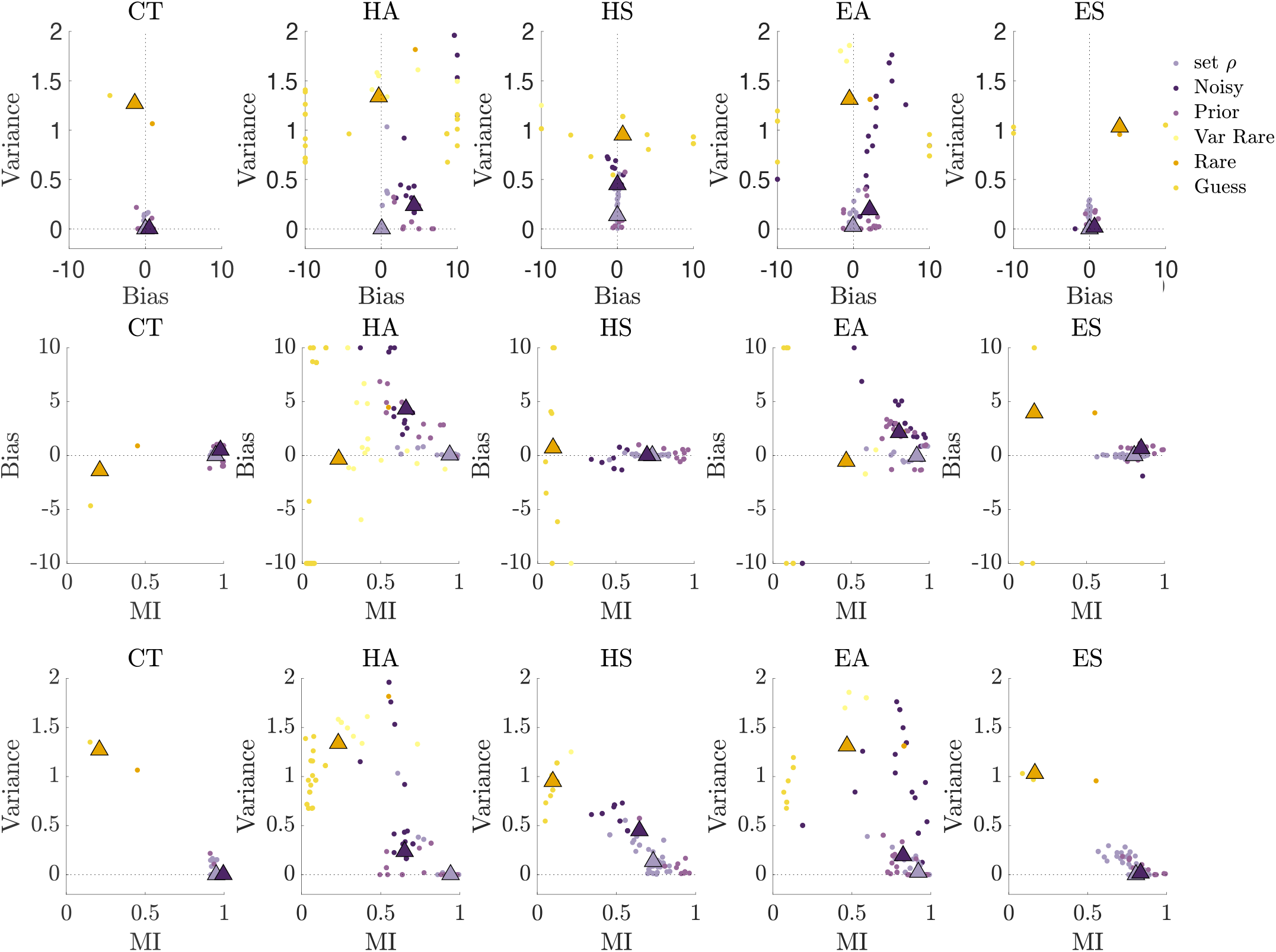
Synthetic data generated using the best-fit model parameters from subjects show representative trends for each model/ class with respect to : (top) bias-variance, (middle) MI-bias, (bottom) MI-variance. Triangles show median heuristic models (yellow), suboptimal Bayesian models (dark purple), and nearly ideal models (light purple). Note that in asymmetric blocks, lower MI Bayesian models show more bias, whereas heuristics show more variance.

**Figure S15:**
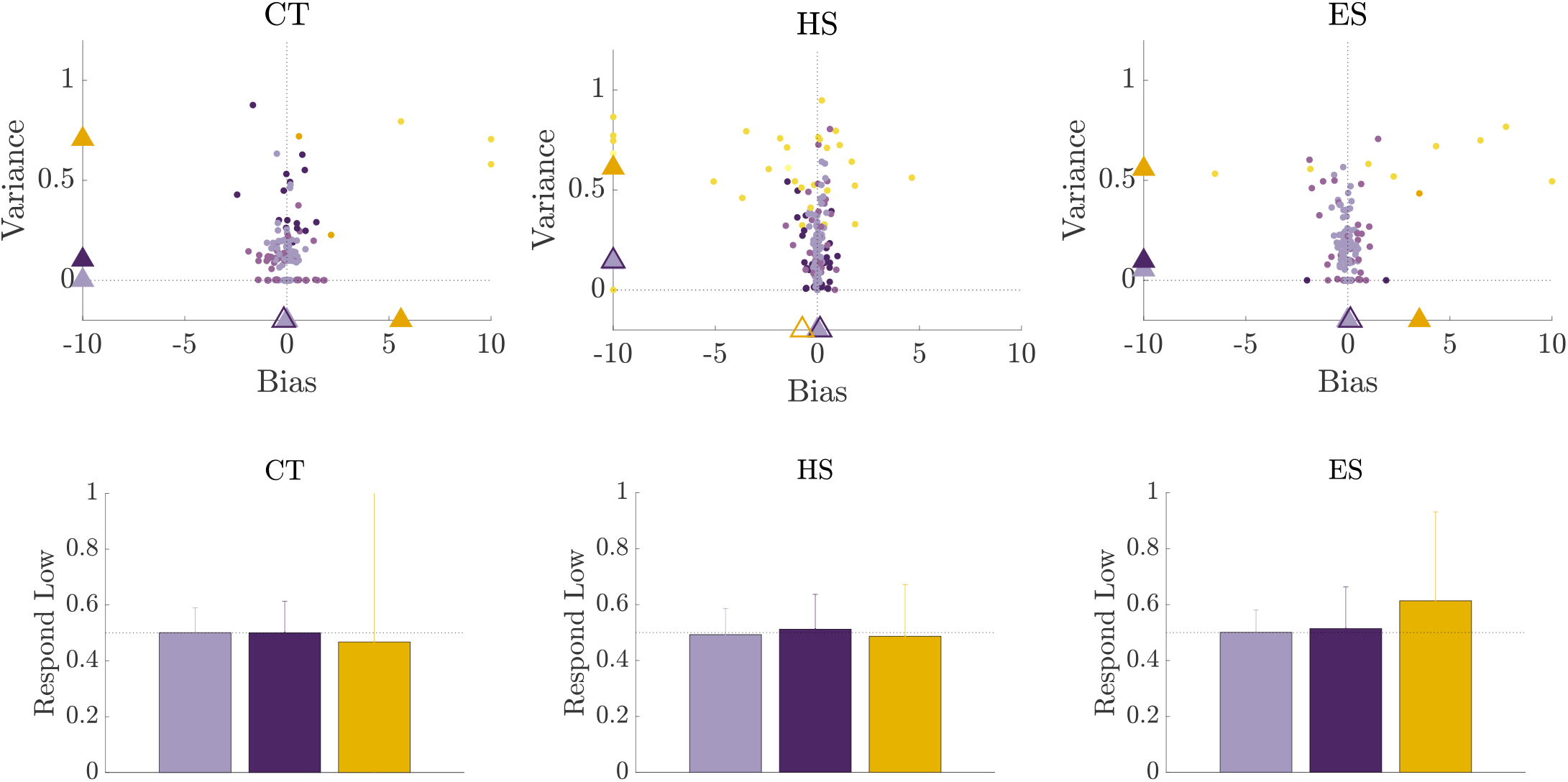
Bias-variance plots for the symmetric task blocks, color-coded by subjects’ best-fit models as in Fig. 3. Triangles represent medians for the the nearly ideal subjects (light purple), suboptimal Bayesian subjects (dark purple), and heuristic subjects (yellow). Filled triangles significantly differ from the nearly ideal subjects based on a two-sided Wilcoxon rank-sum test with *p* < 0.05. Bottom shows corresponding summary of bootstrapped subject low-jar responses (mean and confidence interval) for the nearly ideal subjects (light purple), suboptimal Bayesian subjects (dark purple), and heuristic subjects (yellow).

**Figure S16:**
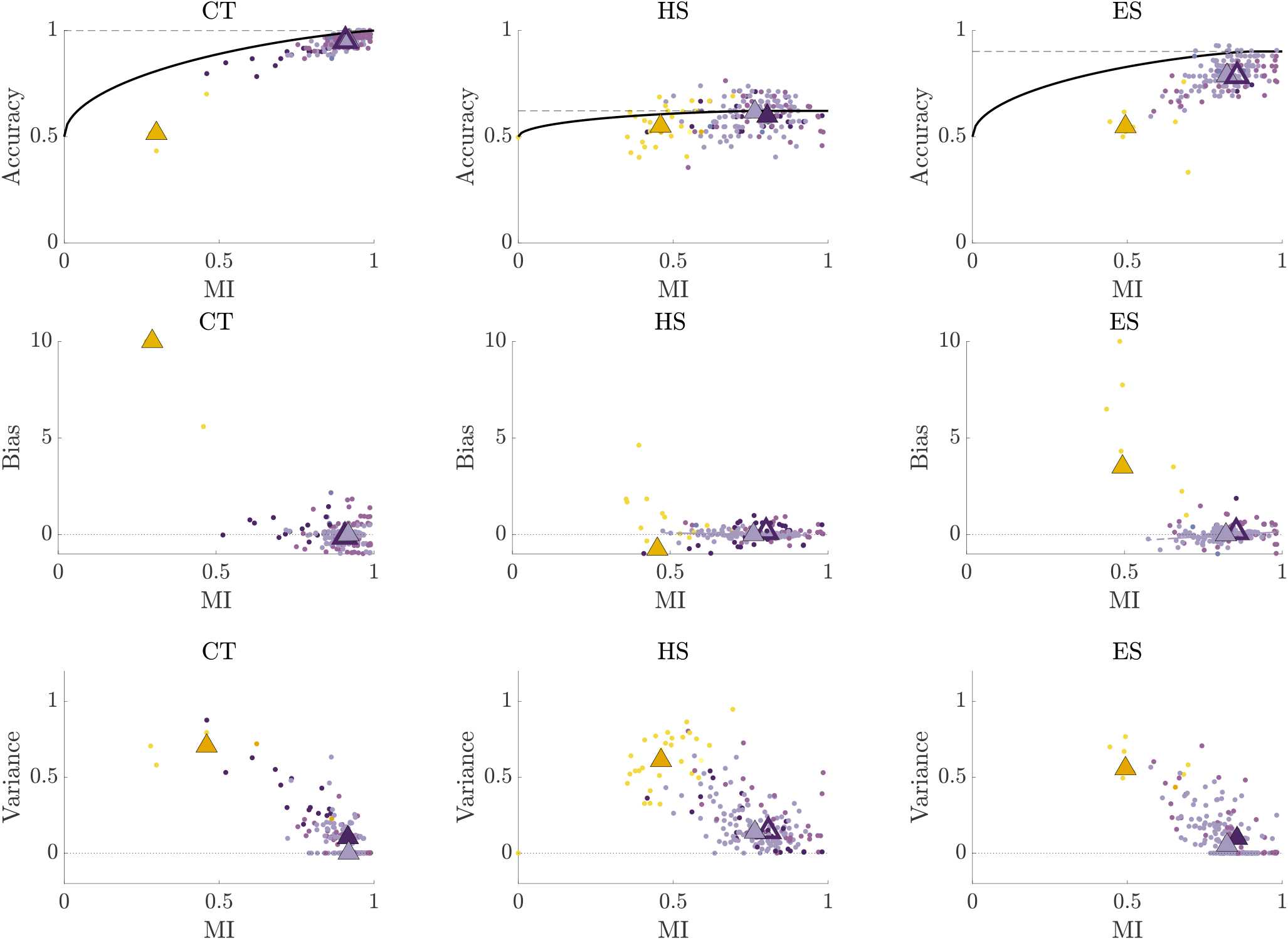
Mutual information and corresponding bias and variance relationships for symmetric blocks, plotted as in Fig. 5**b, c, d**.

**Figure S17:**
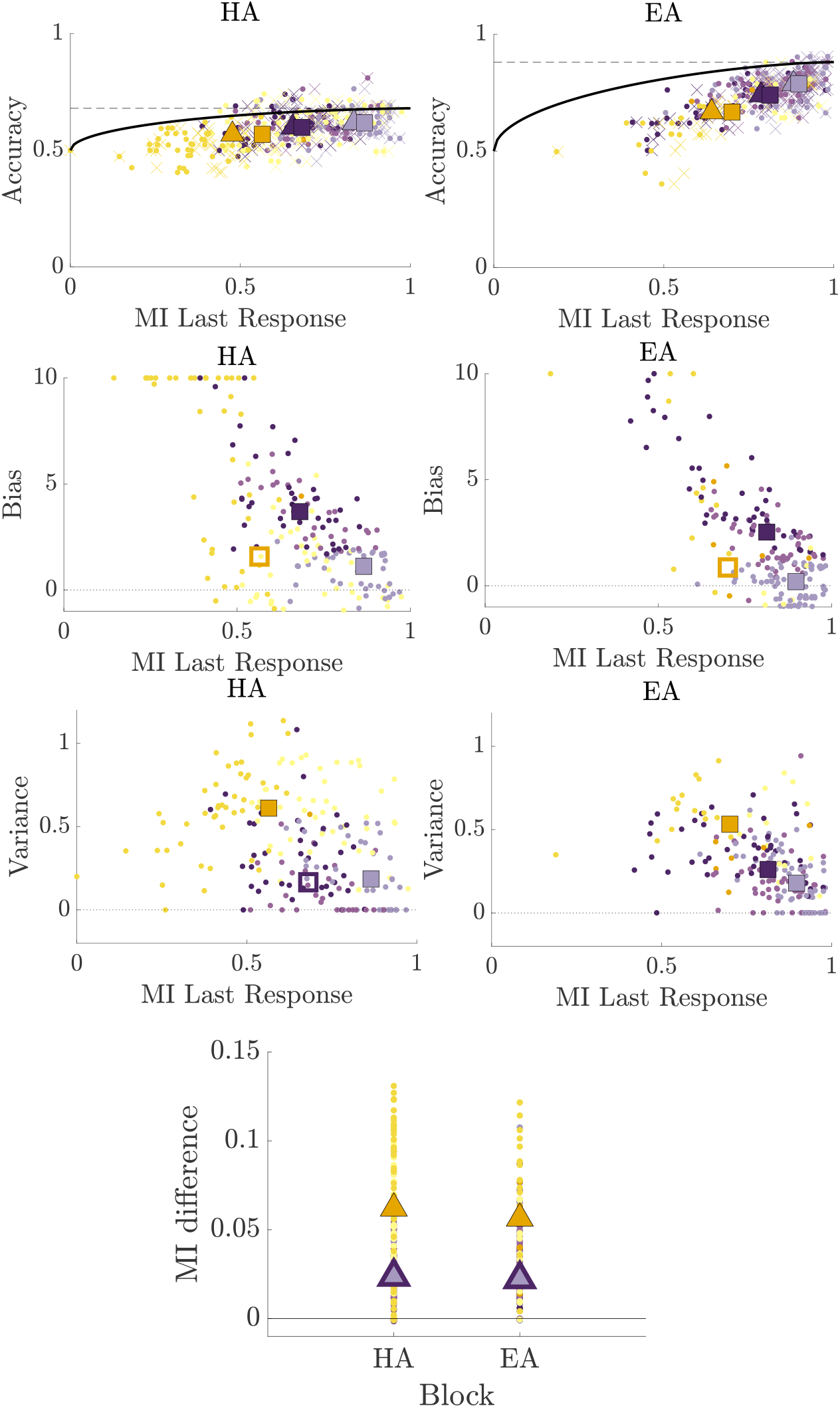
Top: Mutual information including the subject’s response on the previous trial (individual data as xs, medians as squares, significant differences from nearly ideal group are filled) compared to MI without last response (dots, triangles). Middle: MI with last responses and corresponding bias and variance relationships for asymmetric blocks. Bottom plot shows difference in MI with and without previous response for each model group, medians shown as triangles. Significant differences from the nearly ideal subjects shown as solid triangles. In all panels, significance was assessed via a two-sided Wilcoxon rank-sum test with *p* < 0.05. Color codes are as in Fig. 3.

**Figure S18:**
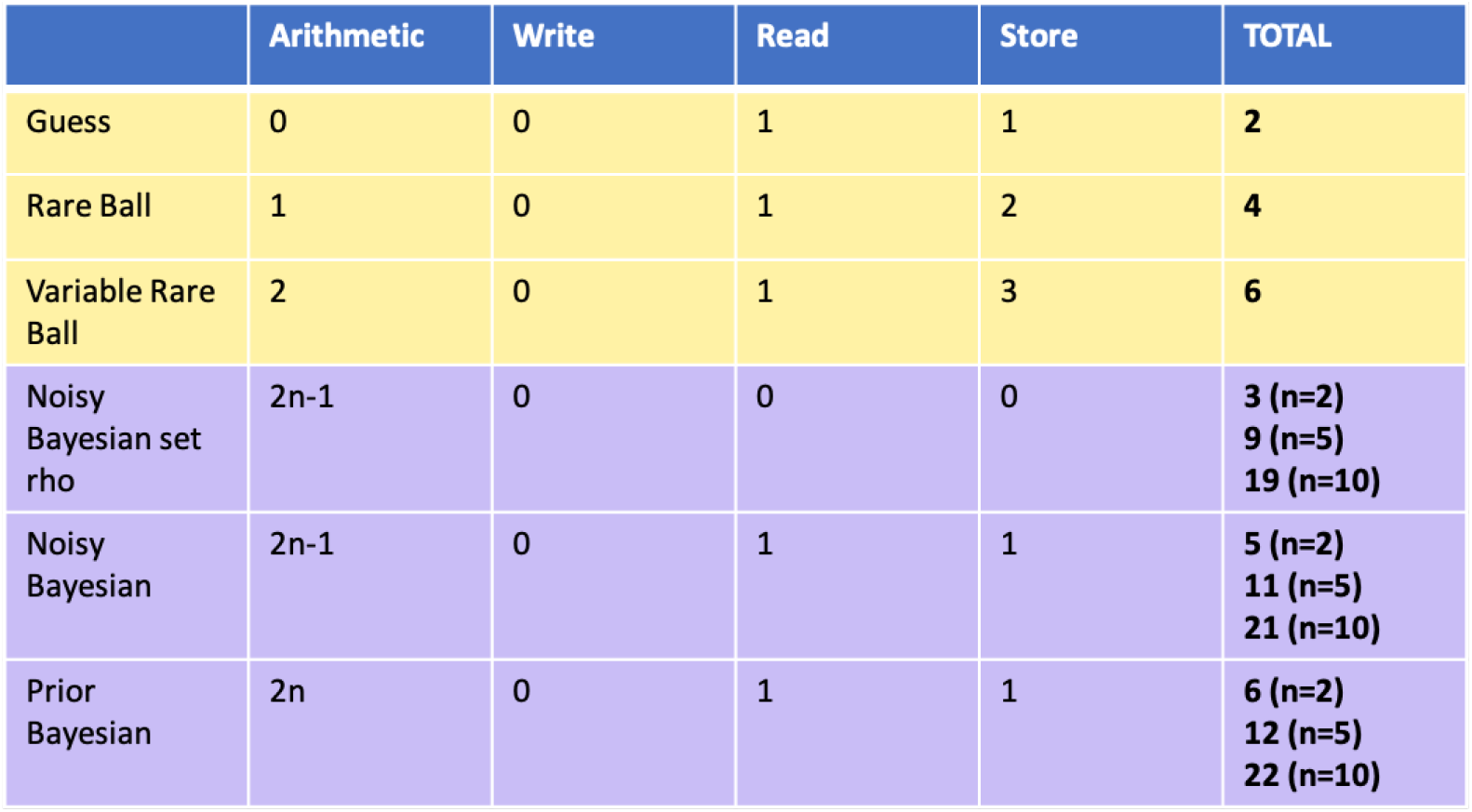
Algorithmic complexity [9] is computed based on the number of operations performed for each trial broken into: arithmetic, writing to memory, reading from memory, and storage. Heuristic models show much less complexity (yellow) compared to Bayesian models (purple). Bayesian model complexity varies with the number of balls observed (*n*). Example computations are shown for sample lengths of 2,5, and 10 balls. Computations are based on the following operations involved in each strategy: **Guess:** Read and store parameter *P*_*guess*_. **Rare Ball:** Identify presence of rare ball (max), read probability of response, store *P*_*rare*_ and *P*_*no*_. **Variable Rare Ball:** All elements from Rare Ball model with additional operations to compute the number of rare balls and store the rare ball threshold *θ*. **Noisy Bayesian Set** *ρ*: Multiplication of the ball weight for each ball observed (*n*) and *n* − 1 summations. **Noisy Bayesian:** Arithmetic as in the Noisy Bayesian Set *ρ* model with additional operations to read and store the rare ball weight *ρ*. **Prior Bayesian** Arithmetic as in the Noisy Bayesian model with inclusion of the prior which is read and stored.

**Figure S19:**
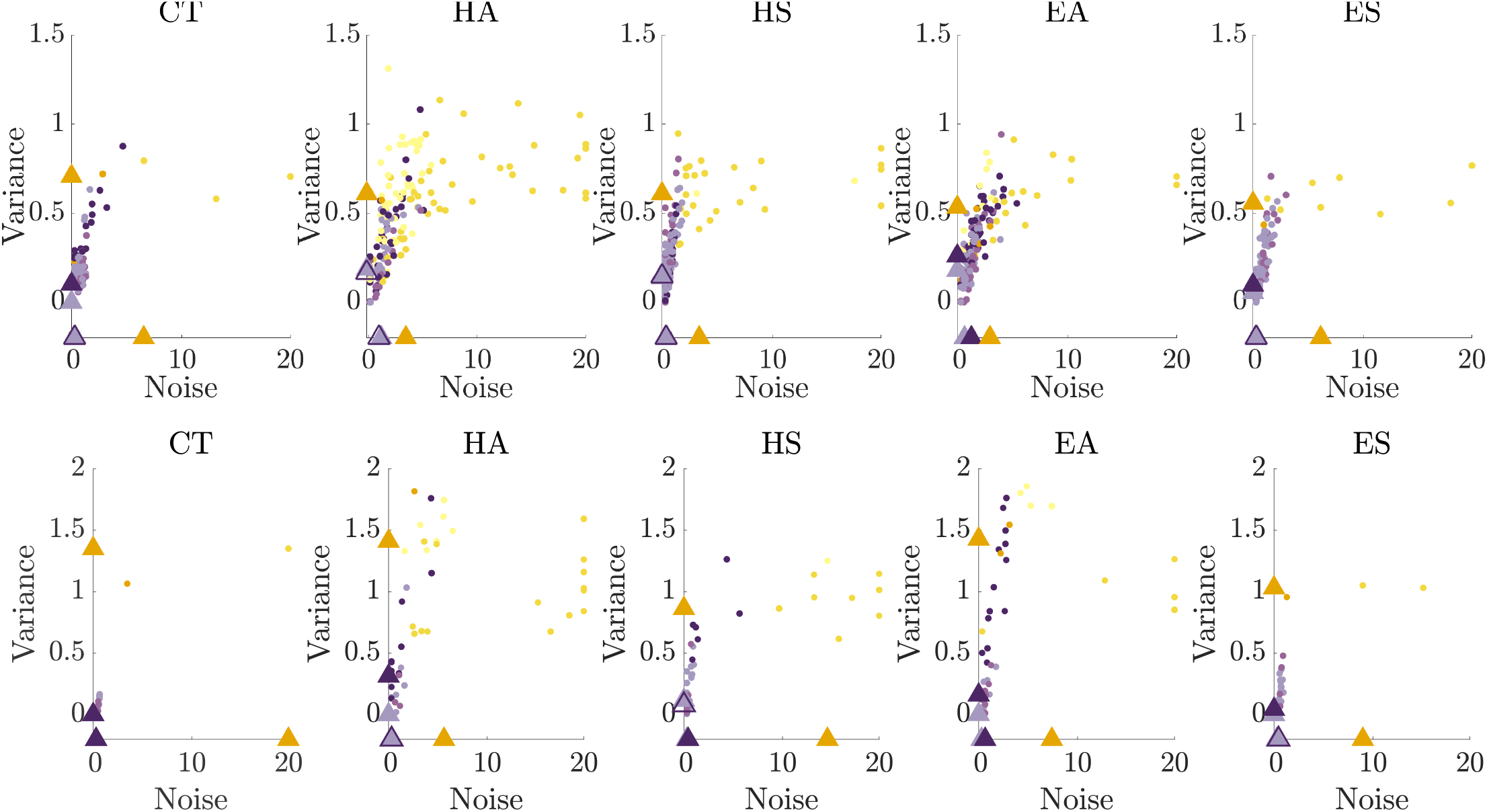
Top: Noise-variance plots for subjects, color coded by the model type that best described a subjects’ responses. Triangles represent medians for the the nearly ideal subjects (light purple), suboptimal Bayesian subjects (dark purple), and heuristic subjects (yellow).Filled triangles significantly differ from the nearly ideal subjects based on a two-sided Wilcoxon rank-sum test with *p* < 0.05 Bottom: Corresponding plots obtained using synthetic data from each model based on subject best-fit parameters. For all plots, large noise values (> 20) were rescaled to 20 for visualization purposes.

**Figure S20:**
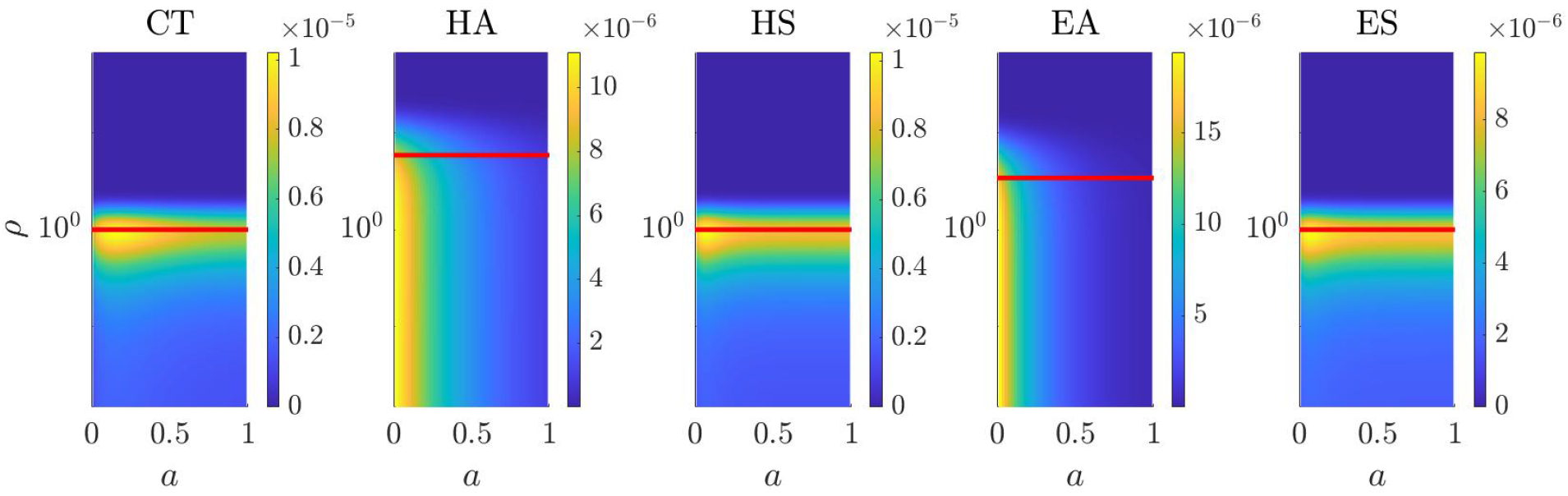
Examples of the weakened informative prior computed from pilot data of 20 subjects. Posteriors were computed for each of the 20 pilot subjects based on the Noisy Bayesian model with a flat prior and then averaged to produce a population posterior for each block. The averaged posterior was then smoothed to create an informative prior (symmetric blocks: L=2, c=5, asymmetric blocks: l=1, c=2). Red line shows the rare ball weighting *ρ* for the ideal observer in each block.

**Figure S21:**
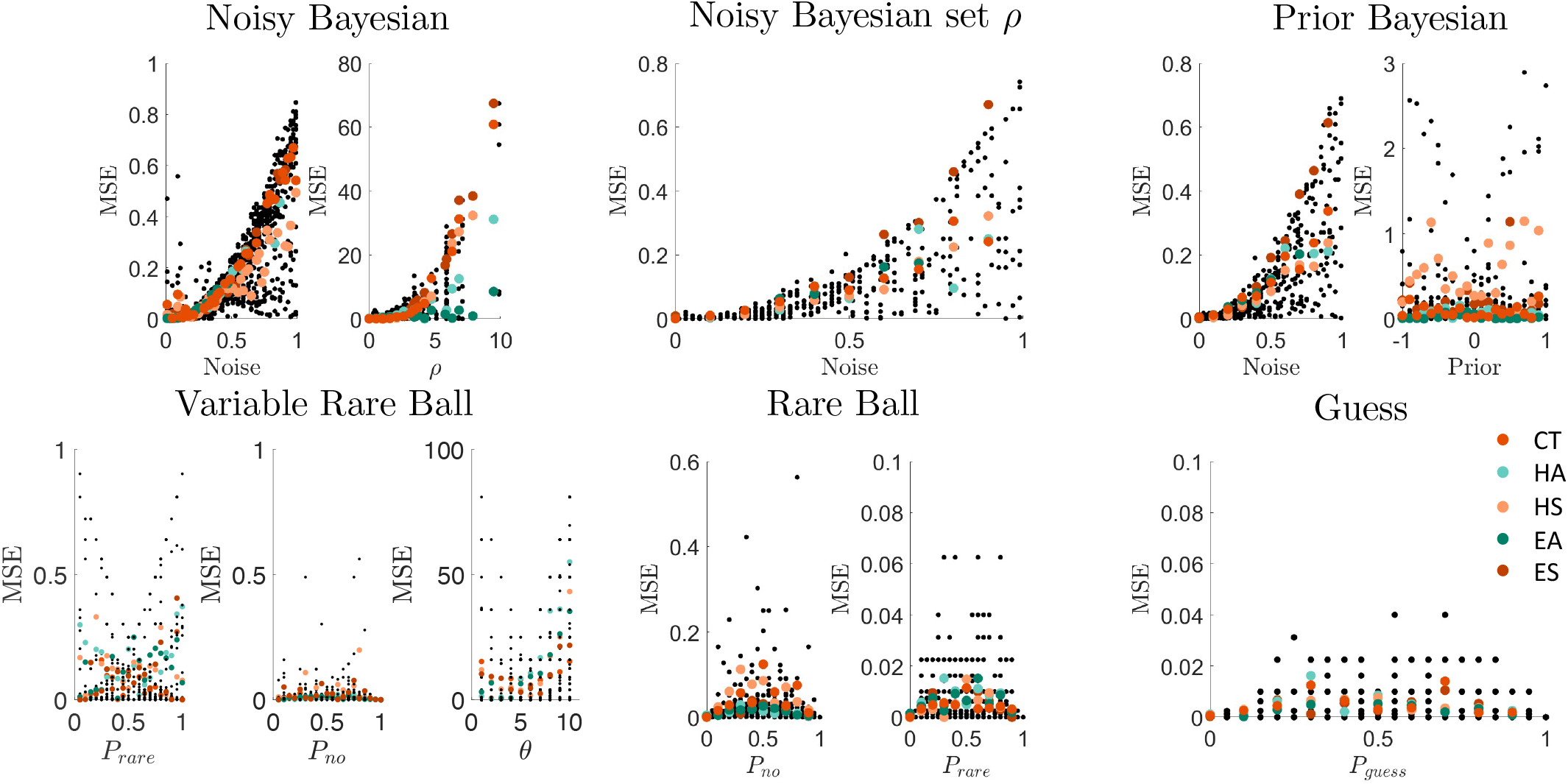
MSE of parameter fits for each model per block based on 100 synthetic datasets with randomly selected parameters using the informative prior developed in Fig. S20 for the Bayesian-based models (top row) and heuristic models (bottom row), as indicated. Black dots represent MSE for synthetic data with a given parameter value (x-axis, other model parameters randomly selected). Colored dots show binned averages for each block. Note that *ρ* is well fit for values near or below the ideal observer’s rare-ball weight, as was seen in the pilot human data. Thus, asymmetric averages (green dots) are closer to zero at higher *ρ* values.

